# Structure of the endocytic adaptor complex reveals the basis for efficient membrane anchoring during clathrin-mediated endocytosis

**DOI:** 10.1101/2020.11.03.364851

**Authors:** Javier Lizarrondo, David P. Klebl, Stephan Niebling, Marc Abella, Martin A. Schroer, Haydyn D.T. Mertens, Katharina Veith, Dmitri I. Svergun, Michal Skruzny, Frank Sobott, Stephen Muench, Maria M. Garcia-Alai

## Abstract

During clathrin-mediated endocytosis, a complex and dynamic network of protein-membrane interactions cooperate to achieve membrane invagination. Throughout this process, middle coat adaptors, Sla2 and Ent1, must remain attached to the plasma membrane to transmit force from the actin cytoskeleton required for successful membrane invagination. Here, we present a cryoEM structure of a 16-mer complex of membrane binding domains from Sla2 and Ent1 that anchors to the plasma membrane. Detailed mutagenesis *in vitro* and *in vivo* of the tetramer interfaces delineate the key interactions for complex formation and deficient cell growth phenotypes demonstrate the biological relevance of these interactions. Finally, time-resolved experiments *in solution* suggest that adaptors have evolved to achieve a fast subsecond timescale assembly in the presence of PIP_2_. Together, these findings provide a molecular understanding of an essential piece for the molecular puzzle of clathrin-coated sites.

## INTRODUCTION

Clathrin-mediated endocytosis (CME) is an essential cellular process that facilitates internalization of external material, integral membrane proteins and lipids into eukaryotic cells. It is involved in many fundamental cellular processes, including nutrient uptake, cell signaling, cell adhesion and polarity, control of plasma membrane homeostasis, and synaptic vesicle recycling. CME also plays an important role in viral infection^1–3^. Clathrin mediates endocytosis through the formation of a protein coat around the endocytic site but it does not interact directly with the plasma membrane. Instead, adaptor proteins such as the AP-2 complex, proteins of the clathrin assembly lymphoid myeloid leukemia (CALM) family and epsins, connect the clathrin coat with the plasma membrane^4–8^. The high-resolution structures of polymerized clathrin coats have provided useful insights into the dynamic endocytic scaffold as a platform for the recruitment of other endocytic proteins and cargo at the membranes of clathrin-coated sites (CCS)^9–12^.

In budding yeast, the endocytic process is well understood, and its timeline and components (with more than 60 proteins involved) have been described in genetic and microscopy studies^2,13–15^. Importantly, though clathrin is present in yeast endocytic sites, it is not absolute prerequisite for endocytosis^16^. In addition, correlative light and electron microscopy (CLEM) has revealed that clathrin does not shape the membrane during invagination, but is instead required to determine the correct size and regularity of the endocytic vesicles^17^. Recent studies suggest that the adaptor proteins epsin Ent1 and Hip1R homolog Sla2 play a critical role in forming the protein coat at yeast endocytic sites^18^.

Adaptors Ent1 and Sla2 co-assemble at the membrane in a phosphatidylinositol 4,5-bisphosphate (PIP_2_)- dependent manner through their epsin N-terminal homology (ENTH) and AP180 N-terminal homology (ANTH) domains, respectively^5,6,18,19^. These adaptors have the topology of elongated knot and string proteins, which connect the plasma membrane and the actin cytoskeleton (through their C-terminal actin-binding domains) allowing them to concentrate the forces provided by actin for successful membrane invagination^18,20–23^. It is known that the PIP_2_ concentration is increased in the plasma membrane at the initial stages of endocytosis promoting the binding of adaptors and mediating the interface of the ANTH-ENTH interaction (from now on abbreviated AENTH) of Sla2 and Ent1^19,24,25^. Yet, the structural role of PIP_2_ during membrane remodeling is still under debate, and our understanding of the nature and dynamics of membrane phospholipids and their associations with endocytic proteins during endocytosis remains limited. It is also intriguing how the weak protein-membrane interactions so far described can hold the plasma membrane during its remodeling without detaching from the membrane^3,5,6^. Another question involving endocytic adaptors is whether the isotropic curvature at the CCS is stabilized by multimeric complex protein scaffolds in addition to hydrophobic insertions of epsin1/2 and AP180 amphipathic α0 helices^5,26–31^. For all these questions the knowledge of the mechanism of ANTH and ENTH recruitment to the PIP_2_-enriched membrane and the detailed structure of AENTH assembly is of the highest importance.

Our study aims to understand the spatial and temporal regulation of the AENTH assembly. We therefore determined the structure of a 16-mer complex of yeast Sla2 ANTH and Ent1 ENTH domains (formed by 8 ANTH and 8 ENTH subunits) bound to PIP_2_ using single-particle cryo-EM and performed stopped-flow Small Angle X-ray Scattering (SAXS) experiments of the complex formation in solution. The structure unveils a hetero-tetrameric assembly of ANTH and ENTH domains as the building unit for larger assemblies. In the AENTH tetramer, the heterodimeric complementary interfaces are mediated by different PIP_2_ molecules clamped between the different domains. Structure-based mutations of these interfaces lead to unstable complexes and growth deficiency phenotypes *in vivo*. The tetramers can be arranged into larger assemblies to adapt to the changing topology of the CCS. We also show that the described ANTH-ENTH interaction is finely tuned as our kinetic studies reveal a fast and efficient self-assembly. *In vitro*, assembly occurs on the hundred millisecond scale and does not require an extra adaptor protein to be present. Finally, we show that the disassembly of the complex is a reversible process that could be simply regulated by decreased local concentrations of any of the partners involved. Altogether, our findings provide a molecular understanding of key adaptor protein structure during endocytic vesicle formation and demonstrate that the mechanism of assembly is not based on weak lipid-protein interactions, but on lipid-mediated oligomeric states.

## RESULTS

### Cryo-EM structure of the AENTH assembly

The ANTH and ENTH domains from Sla2 and Ent1 form discrete protein-lipid complexes in the presence of PIP_2_ in solution^19^. Biophysical characterisation of the sample by SAXS and Dynamic Light Scattering (DLS) revealed a good quality sample of these assemblies (Supplementary **Fig. 1**), which enabled single particle cryo-EM structure determination of a 16-mer complex of ANTH-ENTH with PIP_2_ (**Fig. 1a, 1b**, Supplementary **Fig. 2** and Supplementary **Fig. 3**). The structure reveals an assembly of 8 ANTH and 8 ENTH units arranged into four tetramers, each tetramer consisting of two molecules of ANTH and two of ENTH (A_2_E_2_) (**Fig. 1c** and Supplementary **Fig.4**). The final reconstruction was obtained imposing D2 symmetry **(Fig. 1b** and Supplementary **Fig. 2**). Processing without applying any symmetry produced a similar structure but to lower resolution. The global resolution of the final EM map is 3.9 Å with analysis of the local resolution showing higher resolution towards the core of the map, reaching 3.7 Å (Supplementary **Fig. 3**). Representative images of different regions of the map where the model was built are shown in Supplementary **Fig. 5**. The cryo-EM map shows that each A_2_E_2_ tetramer contains three distinct lipid-binding sites, harbouring 5 lipids per A_2_E_2_ (**Fig. 2**) and a total of 20 PIP_2_ lipids resolved in the entire 16-mer complex. The first binding site is located close to the ANTH KRKH motif, previously reported as a PIP_2_ binding site, involving K24, K26 and H27 and in close proximity to K14 ^5^. Density present at this site could be assigned to the polar head of PIP_2_ used for sample preparation (di-C_8_-PIP(4,5)_2_). Interestingly, the polar head of PIP_2_ seems to also interact with residues K66 and K68 on the ENTH domain adjacent to this site, which together with the ANTH residues, coordinate and keep in place PIP_2_ thus stabilizing the binding of the two domains (**Fig. 2a**). There are 8 PIP_2_ molecules shared in this way between the ANTH and ENTH domains in our 16-mer structure, one per ANTH-ENTH interface. In addition to these eight PIP_2_ molecules, density was present in the previously predicted lipid-binding pocket of the ENTH domain^32^. This binding site involves residues K3 and K10 on the ENTH α0 helix, the amphipathic helix involved in membrane bending, and residues R24, N29, K61 and R62 in proximity to a PIP_2_ molecule (**Fig. 2b**). Each ENTH domain contains one PIP_2_ in its binding pocket, adding 8 PIP_2_ bound to the 16-mer structure. Next to the ENTH PIP_2_ binding site, there is another PIP_2_ molecule (one per A_2_E_2_ tetramer) located in the space between the two α0 helixes of the ENTH domains in between their binding sites. In this case, the polar head of PIP_2_ is coordinated by ENTH K10 and K14 of each of the two ENTH domains, establishing a total of 4 interactions that keep PIP_2_ in place in this interface (**Fig. 2c**). Note that K10 coordinates both the PIP_2_ molecules bound to the ENTH binding pocket and those shared between the two ENTH domains in the tetramer. Each tetramer contains one molecule of PIP_2_ between the two ENTH domains adding a further four in the structure. In summary, our 16-mer A_8_E_8_ structure shows 20 PIP_2_ molecules, all of them clustered close to the core of the structure.

**Fig. 1.**
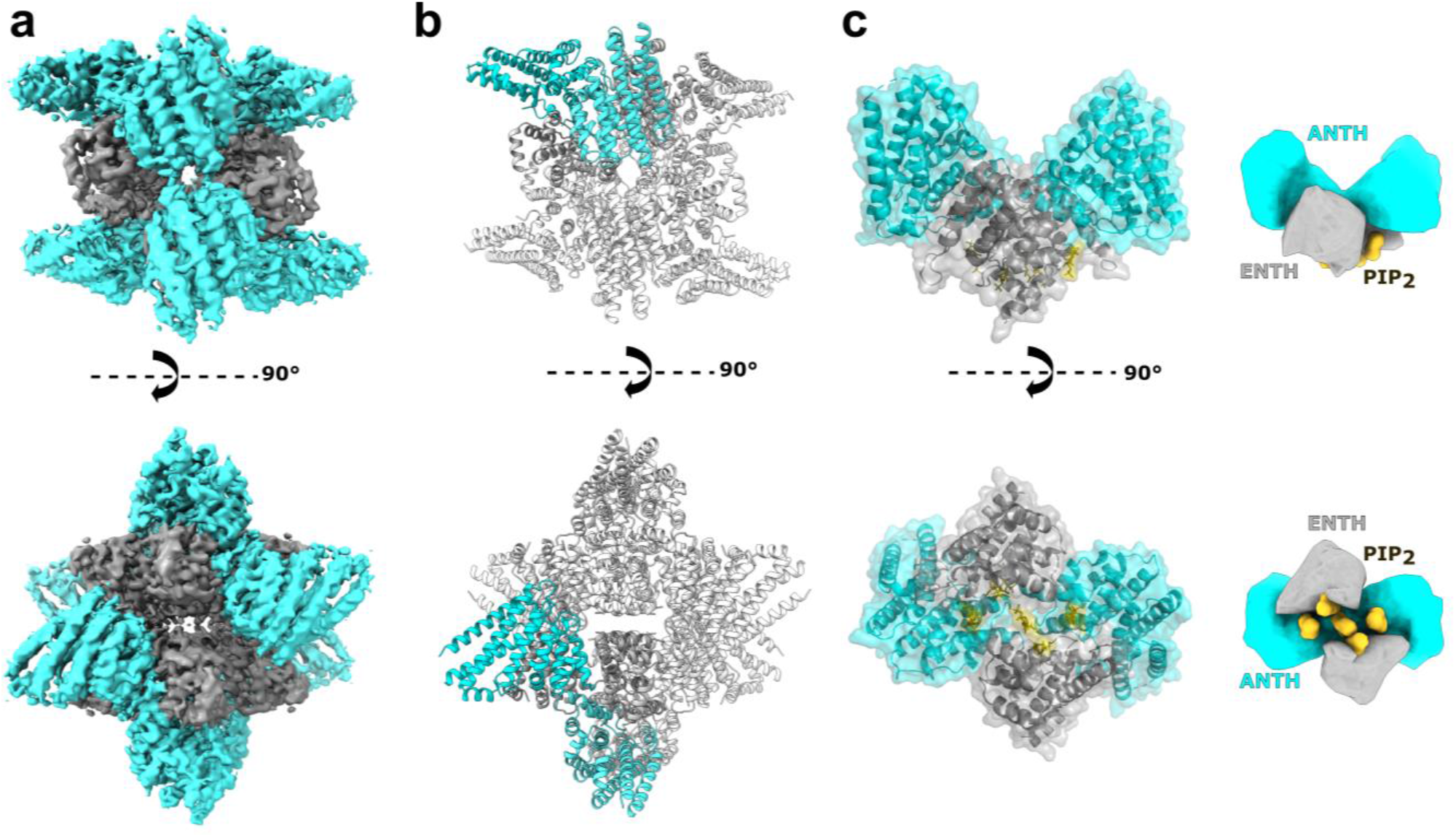
Cryo-EM structure of a 16-mer AENTH complex. **a.** Electron density map obtained for the 16-mer complex formed by 8 ANTH and 8 ENTH units (A_8_E_8_). The density corresponding to ANTH and ENTH is coloured correspondingly in cyan and grey. **b.** Model of the A_8_E_8_ complex built based on the EM density shown in cartoon representation. One tetramer is coloured with the ANTH domains in cyan and ENTH domains in grey (see Supplementary Fig.4 for symmetry axis). **c.** The building unit is the AENTH tetramer (A_2_E_2_) shown in cartoon and surface representation, with the ENTH domains in grey and the ANTH domains in cyan. The PIP_2_ polar heads bound to the tetramer are shown in stick representation (gold). A schematic of the tetramer is shown next to the structure.

**Fig. 2.**
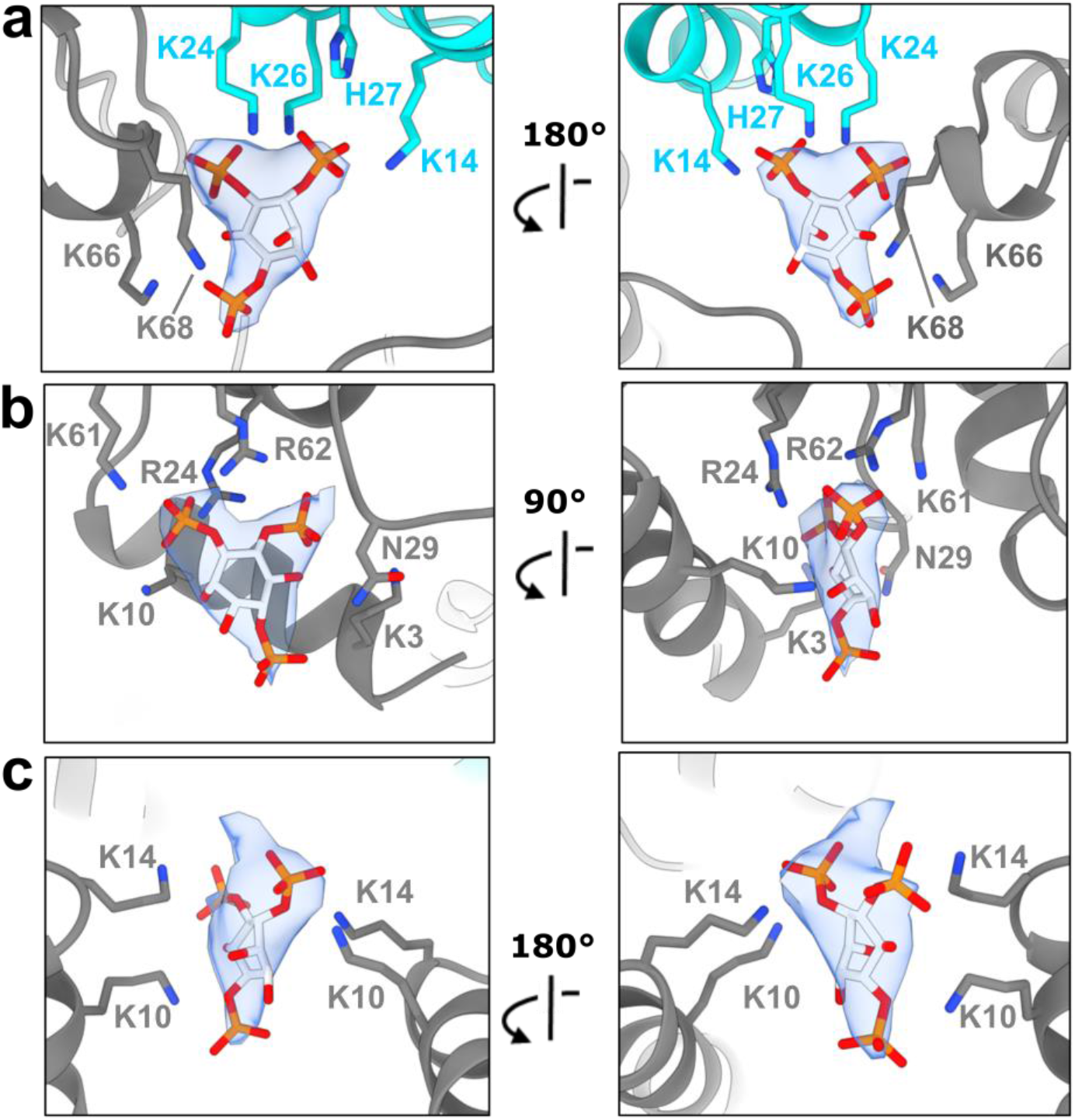
PIP_2_ binding sites in the AENTH tetramer. **a.** PIP_2_ binding site shared by the ANTH and ENTH domains. **b.** PIP_2_ binding site within the ENTH domain. **c.** PIP_2_ binding site in the interface between two ENTH domains. The residues involved in PIP_2_ binding are shown in sticks and coloured in grey for ENTH and cyan for ANTH. The density corresponding to the polar head of the PIP_2_ is shown in mesh.

### The anchor unit of AENTH assembly is a hetero-tetramer

Within each AENTH tetramer, the interactions between the ANTH and ENTH domains are established through two main hetero-dimerization interfaces, and one homo-dimerization ANTH-ANTH interface. Thus, each AENTH tetramer contains a total of 5 protein-protein? interfaces (**Fig. 3**). Previous mutagenesis work on these domains described ENTH T104 and ANTH R29 as important residues for the functionality of Ent1 and Sla2 proteins in yeast ^18,22^. These two residues were found at the centre of an ANTH-ENTH hetero-dimer fitted into a 13.6 Å resolution EM map of the domains’ interaction on GUVs arranged in tubular structures^18^. Our structure shows that ENTH T104 and ANTH R29 are located in one of the new observed interfaces defined as “ANTH-ENTH interface 1” (**Fig. 4a**). Other residues present in this interface are ENTH Y100, E107 and ANTH R25, W36. ANTH R25 is in close contact with ENTH E107 and ENTH Y100 is oriented parallel to ANTH W36, establishing a stacking interaction by the coordination of their aromatic rings (**Fig. 4a**). To test the relevance of this interface for complex formation, point mutations were introduced in residues in the ANTH and ENTH domains present in this interface (Supplementary **Fig.6a**). AENTH complexes were assembled in the presence of PIP_2_ *in vitro* as done for the cryo-EM sample preparation using mutants for one of the domains and its wild-type counterpart to assess the impact on the complex assembly. Wild type ANTH and ENTH domains in the presence of PIP_2_ form structures around 8 to 10 nm of average hydrodynamic radius in DLS experiments. Mutation of these residues showed less particles corresponding to 16-mer assemblies *in vitro* when compared with wild-type complexes **(Fig. 4d and Table 2)**. Non-denaturing electrospray ionization mass spectrometry (native MS) of the selected mutant ENTH Y100R showed mostly monomeric species instead of 16-mer assemblies (**Fig. 4g**), thus confirming the disruption of the complex. The phenotype of Y100R mutant was previously attributed to a possible role of Ent1 in binding GTP-activating proteins for Cdc42, a critical regulator of cell polarity ^33^. Here, we show that Y100 is key for the AENTH complex formation. In addition, mutation *in vivo* of ANTH R25, R29 and W36 and ENTH Y100 and E107 showed strong or intermediate growth deficiency phenotypes in yeast, confirming an essential role of this interface for proper endocytic function (**Fig. 4h and 4i**). Mutation of ENTH residue F108 also introduced a growth defect phenotype *in vivo* (**Fig. 4i**), but our *in vitro* data showed that the stability of the protein is compromised by this mutation (Supplementary **Fig. 6**). A similar scenario was found for ANTH Y247 and L248, conserved specifically in ANTH domains of the Sla2 family, whose deletion caused a endocytosis-linked growth defect in yeast, previously attributed to a possible interface in the protein complex with epsin^19^. Our structure shows for the first time that these residues are not involved in any interface of the assembly, but instead that their deletion leads to an unstable protein *in vitro* suggesting that they are crucial for the folding of ANTH (Supplementary **Fig. 7**).

**Fig. 3.**
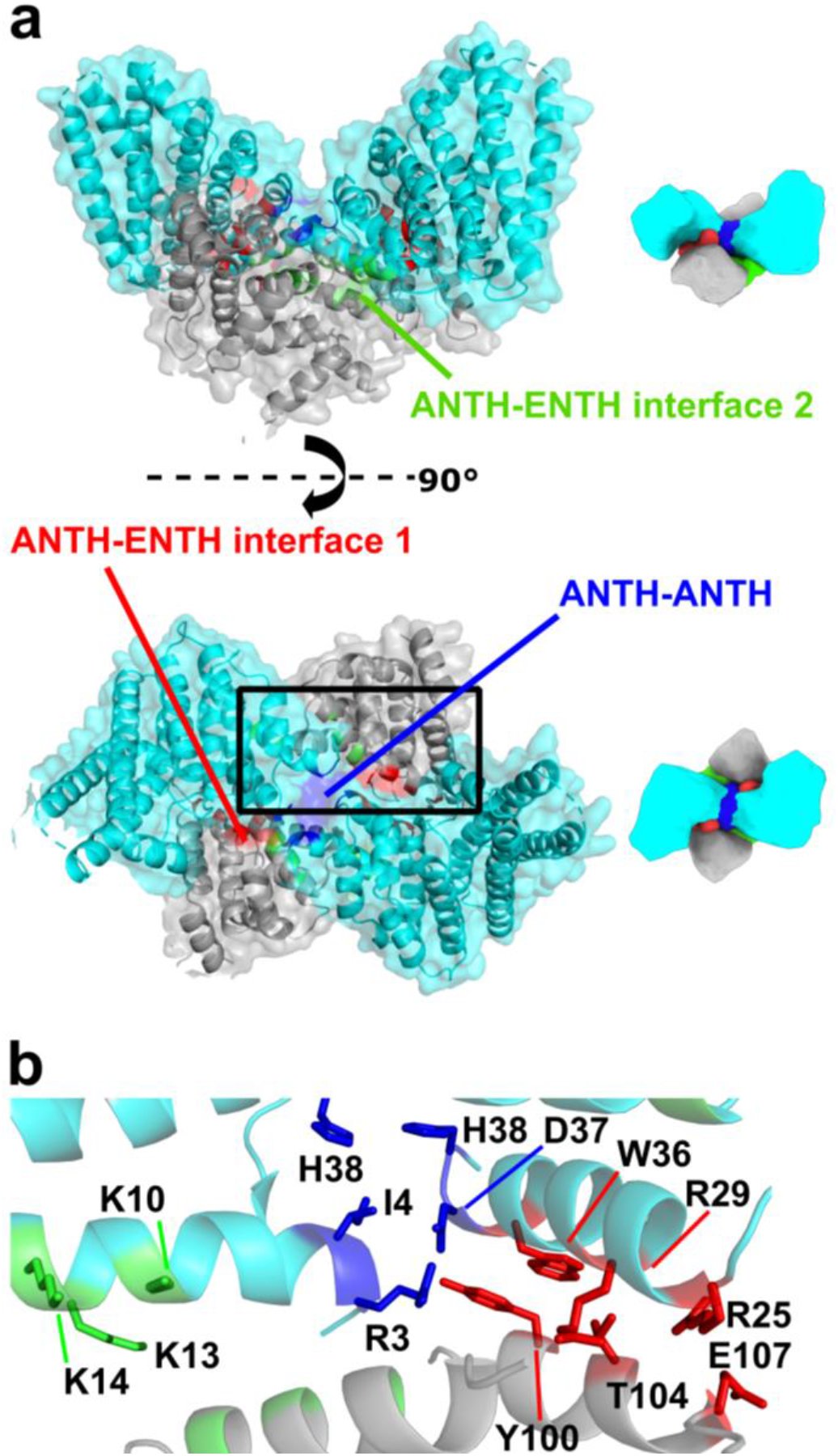
The AENTH tetramer interfaces. **a.** Surface representation of the A_2_E_2_ tetramer, with ANTH subunits in cyan and ENTH subunits in grey. The different interfaces are coloured: ANTH-ENTH interface 1 in red, ANTH-ENTH interface 2 in green and ANTH-ANTH interface in blue. **b.** Cartoon representation of the tetramer area within the black rectangle in a showing the different interface residues involved in sticks (same color code as in **a**).

**Fig. 4.**
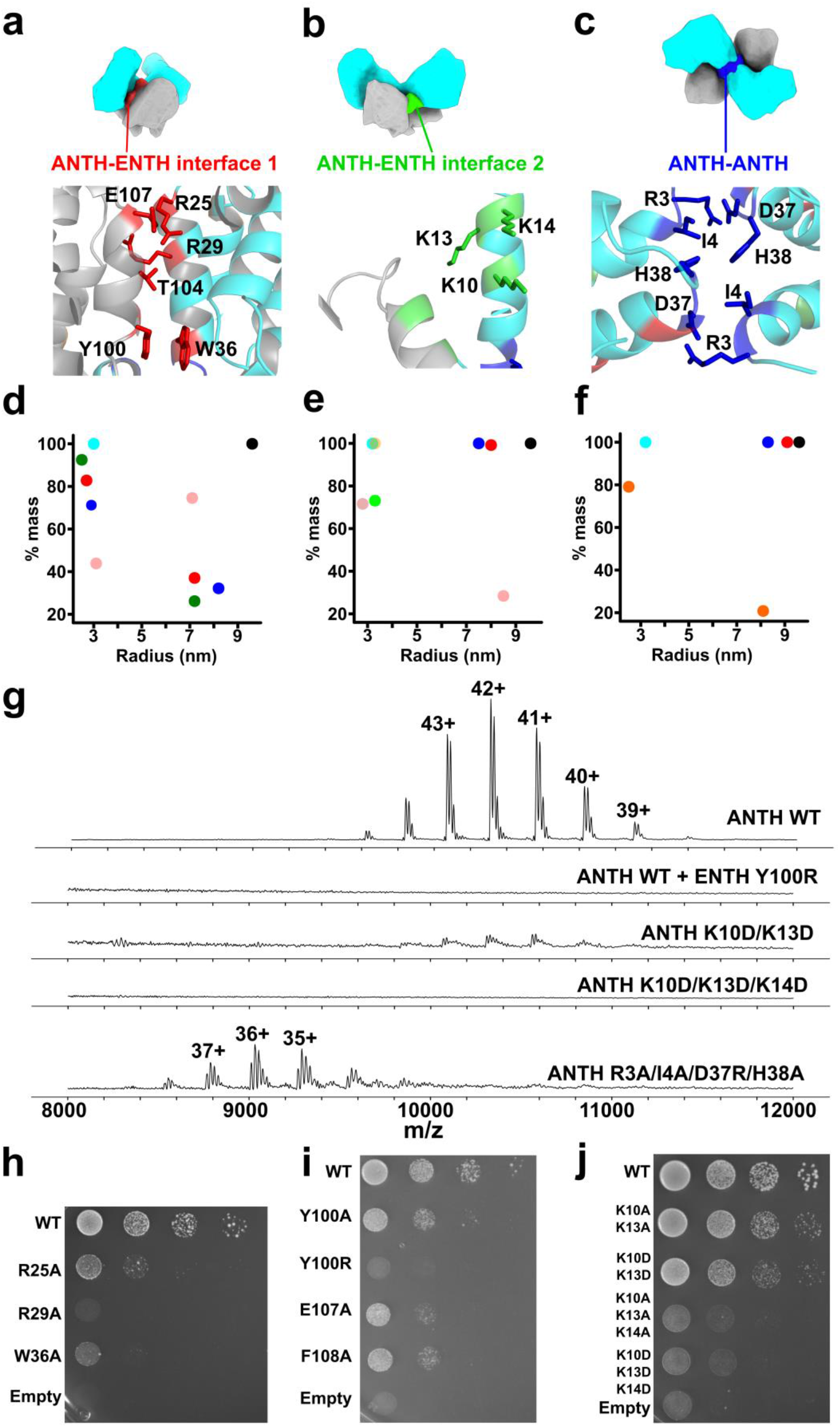
Different interfaces present in the AENTH tetramer. **a-c.** Schematic representation of the AENTH model showing the different interfaces present, designated as ENTH-ANTH interface 1 (**a**), ENTH-ANTH interface 2 (**b**) and ANTH-ANTH-interface (**c**). In the lower panels, the residues involved in each of the interfaces are shown in stick representation and coloured using the color scheme for each of the interfaces. **d-e.** Intensity mass plot from DLS data showing the hydrodynamic radius of particles in solution for wild type AENTH (black) and different mutants. As a reference the ANTH domain is shown in cyan. **(d)** ENTH-ANTH interface 1 showing mutations ANTH Y100R (red) ANTH R25A (blue), ANTH R29A (green) and ENTH E107A (pink); **(e)** ENTH-ANTH interface 2 with mutations ANTH K10A/K13A/K14A (pink) ANTH K10D/K13D/K14D (yellow), ENTH E54A/D57A/D60A (blue), ENTH E54A/D57A (green) and ANTH Q9A/K10A (red). (**f**) ANTH-ANTH interface with mutations ANTH R3A (orange), ANTH R3A/I4A/D37R/H38A (blue) and ANTH R3A/I4A/D37R (red). **g.** Summary of the native mass spectrometry results obtained for the mutants of the tetramer interfaces. **h-j.** Growth defects of mutants of ANTH (**h**) and ENTH (**i**) ANTH-ENTH interface 1, and ANTH-ENTH interface 2. **(j)**. Interface mutants were expressed after depletion or deletion of endogenous Ent1 and Sla2 proteins, respectively. Cell growth was analyzed by plating 10-fold serial dilution of cells on SD-Ura plates and incubated for 3 days at 37 °C.

**Table 1:** Cryo-EM data collection, refinement and validation statistics of the AENTH 16-mer assembly

|  |  |
| --- | --- |
| **Data collection and processing** |  |
| Magnification | ×75,000 |
| Voltage (kV) | 300 |
| Electron exposure (e<sup>-</sup>/Å<sup>2</sup>) | 75.2 |
| Defocus range (μm) | −1 to −3 |
| Pixel size (Å) | 1.065 |
| Symmetry imposed | D<sub>2</sub> |
| Initial particle images (no.) | 195,536 |
| Final particle images (no.) | 79,414 |
| Map resolution (Å) | 3.9 |
| FSC threshold | 0.143 |
| Map resolution range (Å) | 3.7 to 5.7 |
| **Refinement** |  |
| Initial model used | 5onf, 5oo7 |
| Model resolution (Å) | 3.9 |
| FSC threshold | 0.5 |
| Model resolution range (Å) | - |
| Map sharpening *B* factor (Å<sup>2</sup>) | −200 |
| **Model composition** |  |
| Non-hydrogen atoms | 26,192 |
| Protein residues | 3152 |
| Ligands | 20 |
| ***B* factors (Å<sup>2</sup>)** |  |
| Protein | 70 |
| Ligand | - |
| **r.m.s. deviations** |  |
| Bond lengths (Å) | 0.012 |
| Bond angles (°) | 2.037 |
| **Validation** |  |
| MolProbity score | 1.91 |
| Clashscore | 2.60 |
| Poor rotamers (%) | 3.78 |
| **Ramachandran plot** |  |
| Favoured (%) | 93.56 |
| Allowed (%) | 5.80 |
| Disallowed (%) | 0.64 |

**Table 2.** ANTH-ENTH complexes measured by DLS

| **Mutant complex** | **Radius**<br>**(nm)** | **%Pd** | **Mw-R**<br>**(kDa)** | **%Intensity** | **%Mass** | **%Number** |
| --- | --- | --- | --- | --- | --- | --- |
| **ANTH M1A-ENTH WT** | 3.3 | 15.49 | 53.2 | 10.8 | 69.5 | 96.7 |
|  | 8.5 | 20.4 | 501.4 | 71.4 | 30 | 3.3 |
| **ANTH R3A/I4A/D37A-**<br>**ENTH WT** | 8.3 | 9.15 | 479.8 | 100 | 100 | 100 |
| **ANTH S6A/K10A-ENTH**<br>**WT** | 6.7 | 6.86 | 287.7 | 93.8 | 99.4 | 100 |
| **ANTH K13A-ENTH WT** | 7.2 | 10.19 | 344.6 | 100 | 100 | 100 |
| **ANTH A15A-ENTH WT** | 6.2 | 15.99 | 236.5 | 99.4 | 100 | 100 |
| **ANTH Q9A/K10A-**<br>**ENTH WT** | 8 | 11.18 | 433.9 | 96.8 | 99.2 | 100 |
| **ANTH WT-ENTH**<br>**F5A/I12A** | 8 | 11.62 | 436 | 100 | 100 | 100 |
| **ANTH WT-ENTH**<br>**F52A/F53A** | 7.9 | 11.44 | 425.5 | 95.7 | 99.5 | 100 |
| **ANTH WT-ENTH**<br>**E54A/D57A** | 7.5 | 26.06 | 374 | 100 | 100 | 100 |
| **ANTH WT-ENTH Y100R** | 2.7 | 0 | 33.3 | 15.2 | 78.5 | 98.1 |
|  | 7.2 | 14.81 | 344.4 | 71.3 | 21.3 | 1.9 |
| **ANTH WT-ENTH WT** | 9.6 | 11.85 | 664.6 | 100 | 100 | 0 |
| **ANTHK10A/K13A/K14A-**<br>**ENTH WT** | 8.5 | 16.31 | 508 | 90.7 | 28.4 | 1.6 |
| **ANTHΔY247/L248-**<br>**ENTH WT** | 9 | 36.27 | 571.8 | 100 | 100 | 100 |
| **ANTH**<br>**K10D/K13D/K14D-ENTH**<br>**WT** | 3.2 | 12.67 | 51.4 | 39.8 | 99.9 | 100 |
| **ANTH WT-ENTH Q20A** | 8.6 | 11.95 | 517 | 97.2 | 98.8 | 100 |
| **ANTH WT-ENTH E97A** | 9.9 | 23.66 | 727.2 | 99.2 | 56.3 | 1.9 |

A second interface in the AENTH tetramer involves residues K10, K13 and K14 of ANTH (**Fig. 4b** Supplementary **Fig.6b**). ANTH K14 is involved in the interaction with PIP_2_, indicating synergy in PIP_2_ binding and function of this ANTH-ENTH interface 2. To address the importance of this interface, point mutations were introduced in this interface. After testing that mutations did not affect stability of individual domain (**see Methods**), we observed that mutants from the ANTH-ENTH interface 2 display complexes with a lower aggregation temperature (*T_agg_*) by nanoDSF, indicating that the stability of these complexes is partially compromised (Supplementary **Fig. 6e**). Apart from this destabilization effect, some mutants also showed a clear effect on complex assembly reported by DLS (**Fig. 4e and Table 2**) when compared with wild type AENTH complex. Specifically, triple mutant ANTH K10A/K13A/K14A showed a smaller proportion of 16-mer assemblies in solution, while mutant ANTH K10D/K13D/K14D completely abolished complex formation **(Fig. 4e).**

To explore with a greater level of detail the effect of these mutations for the assembly of the ANTH and ENTH domains, native MS was used to analyse the complexes obtained when mutant domains were used to reconstitute the assembly. ANTH K10D/K13D displays a large destabilization of the complex for this interface observed by nanoDSF (Supplementary **Fig. 6e**) and only a weak Native MS signal corresponding to the 16-mer assembly (**Fig. 4g**), when compared to the wild-type. Both results indicate that K10 and K13 are important for oligomerization. Beyond that, ANTH K10D/K13D/K14D completely abolished complex formation, and no signal for larger assemblies other than monomeric species was observed by native MS in agreement with the DLS data (**Fig. 4g**). These results indicate that the interactions in the interface 2 are essential for proper complex assembly. *In vivo*, mutations of these residues caused growth defect phenotypes in yeast strains lacking Sla2 (**Fig. 4j** and **Supplementary Fig. 8b**). Mutation of ANTH K10/K13 to alanine caused an intermediate growth defect phenotype enhanced when these residues were mutated to glutamic acid. The tripe mutant K10/K13/K14 present in this interface caused a severe growth defect phenotype when mutated to alanine, further enhanced when replaced by glutamic acid. Furthermore, each of the three lysine residues has an important role in complex formation, as individual mutations of these lysines also caused growth defect phenotypes *in vivo* (Supplementary **Fig. 8b**). Altogether, these results indicate that lysines of the ANTH-ENTH interface 2 are essential for the assembly of the tetrameric lipid-binding AENTH unit and for the AENTH function in vivo. Interestingly, on the ENTH domain, E54A/D57A/D60A showed the largest destabilization in the nanoDSF data (Supplementary **Fig. S6e**). However, mutation of these residues did not show any complex disruption by DLS *in vitro* nor introduced a growth defect in yeast cells, most likely ruling out a major role in the AENTH assembly and function (Supplementary **Fig. 8a**).

Finally, ANTH-ANTH interface mutants (Supplementary **Fig. 6c**) did not show a large destabilization effect over the complex *in vitro* with the exception of ANTH R3A (Supplementary **Fig. 6f**) which also generated a larger amount of monomeric species upon complex formation when compared to the wild type (**Fig. 4f and Table 2**). Interestingly, native MS for the R3A/I4A/D37R/H38A mutant showed a shift in the signal of the complexes obtained to lower masses, in agreement with the DLS data (**Fig 4g**). However, this mutation did not cause severe growth defect phenotype *in vivo* (Supplementary **Fig. 8c**).

### The AENTH tetramer can form larger assemblies

Previous work showed that the ANTH and ENTH domains together tubulate GUVs and coat them with regular helical assemblies, as determined by cryo-EM to 13.6 Å resolution^18^. Our 16-mer structure solved by single particle EM shows that, instead of the heterodimer as previously proposed, the AENTH tetramer is the building unit of larger clusters and assemblies. Therefore, we used our AENTH tetramer for flexible fitting into the electron density map of ANTH-ENTH coat on GUVs using adaptive distance restraints in ISOLDE (Supplementary **Fig. 9**). Overall, the fit agrees with the previous assignment of the domains to the larger and smaller densities present on the surface of the tubules for the ANTH and ENTH domains respectively. It also places the ENTH α0 helixes pointing towards the core of the tubules (Supplementary **Fig. 9a**). This is also consistent with the membrane bending mechanism of the ENTH domain by insertion of the α0 helix and displacement of the lipids in the inner layer of the plasma membrane^35^. Similarly, the fit also shows that the ANTH domain has its lysine patch available to bind the polar heads of the PIP_2_ at the membrane, in agreement with its suggested lipid binding mechanism of these two domains. In spite of the close similarity between the membrane bound model and the structure in solution presented here, the distance between the two ENTH domains of the tetramer seem to be much larger in the membrane-bound model. Therefore, the membrane bound tetramer is in a more “open” conformation possibly as a consequence of the presence of a membrane, which causes the tetramer to “open” compared to the structure solved with PIP_2_ in solution (Supplementary **Fig. 9b**). Mechanistically, this difference between the membrane-bound and the solution structure could potentially reflect that the tetramer exerting some force on the membrane by clamping of the ENTH domains.

In our 16-mer assembly, all the PIP_2_ polar head groups are located close to the core of the structure (**Fig. 5a**). Since samples were prepared above the critical micelle concentration (CMC) of di-C_8_-PIP_2_^36,37^, the position of the polar groups indicates that the core of the map is composed of a PIP_2_ micelle around which the ANTH and ENTH domains assemble. Nevertheless, there is no density present at the core of the map, where possibly the hydrocarbon tails of the lipids might be present but their flexibility could therefore prevent them from being resolved. PIP_2_ micelles were measured during our DLS experiments and showed a hydrodynamic radius around 0.9-1.4 nm. In addition, all ENTH domains arrange themselves around the micelle displaying their α0 helixes into its hydrophobic core (**Fig. 5b**). All this suggests that our 16-mer AENTH complex is assembled on a minimal micelle made of di-C8-PIP2 molecules.

**Fig. 5.**
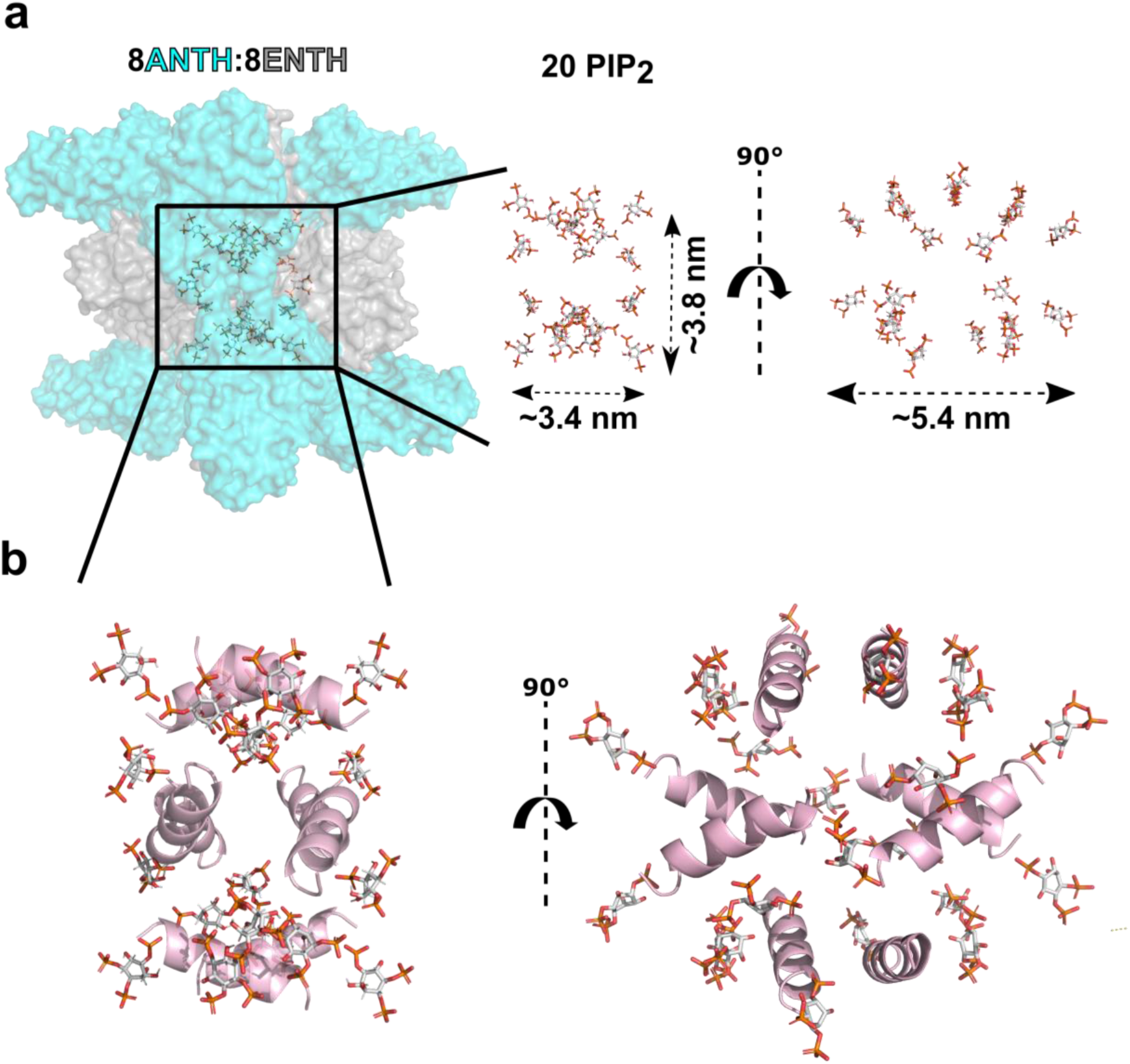
Details of the core of the AENTH 16-mer complex structure. **a.** All the polar heads from the PIP_2_ placed in the structure (shown in sticks) are contained in the region near to the core of the structure (shown in surface representation), indicating the presence of a PIP_2_ micelle in the center of our map. **b.** Only the α0 helices from ENTH subunits are shown in cartoon representation (in pink) together with the PIP_2_ shown in stick, all of them are pointing towards the interior of the structure.

Interestingly, under the experimental conditions used during our Native MS experiments, ANTH and ENTH domains form 16-mer assemblies as determined by cryo-EM (A_8_E_8_). Additional assemblies have been previously described for AENTH corresponding to 6 ANTH and 6 ENTH molecules (termed A_6_E_6_)^19,34^. We observed this A_6_E_6_ assembly was observed for some ANTH-ANTH interface mutants and for the ENTH α0 helix mutant F5A/L12A/V13A, which produced a native MS spectrum with signal only corresponding to the 12-mer assembly, and was later confirmed by cryoEM (Supplementary **Fig. 10** and Supplementary **Table 1**). The map obtained for the assembly of this mutant showed density which could be unambiguously assigned to three A_2_E_2_ tetramers arranged around a central core of PIP_2_ in a similar fashion as the A_8_E_8_ assembly (Supplementary **Fig. 10b** and Supplementary **Fig. 10c**). The position of the ANTH and ENTH domains is remarkably similar within each hetero-tetramer (A_2_E_2_), indicating that mutation of hydrophobic residues on the amphipathic helix does not disrupt the ability of the ENTH domains to assemble into the functional hetero-tetramer with the ANTH domains. All these observations support the hypothesis that the building units of ANTH complexes (either in solution or on GUV membranes) are hetero-tetramers that build the different assemblies.

### Kinetics of the AENTH complex assembly

In equilibrium, ENTH and ANTH form protein-lipid clusters in the presence of PIP_2_, whose major component is the A_8_E_8_ AENTH complex determined by cryo-EM. Supplementary **Fig. 1** shows the SAXS curve for that equilibrium in solution. In order to study compatibility with cellular time scales of such assembly, we performed stopped-flow SAXS (SF-TR-SAXS) studies of the interaction upon fast mixing of its constituents. A stopped-flow device enabled rapid mixing of the ANTH and ENTH domains followed by immediate exposure to X-ray radiation and the scattering signal followed over time. The buffer-subtracted SAXS curves obtained over the first 400 ms are shown in **Fig. 6a**. We observed a very fast change of the SAXS curves, particularly in the q-region between 0.02 to 0.6 nm^−1^ over this period of time, after which the curves remained relatively unchanged. The fitting of the early points of the assembly (up to 300 ms) to a single exponential binding equation gave a first exponential kinetic constant of around ≈ 90 ms (**Fig. 6b)**, showing a fast increase in the contribution of the component of the data accounting for the complex formation in solution. In conclusion, our SF-TR-SAXS data indicate that the association of the adaptors takes place quickly and equilibrates mainly to the 16-mer assembly in solution. The kinetic constants obtained in solution are thus in agreement with the fast oligomerization most probably occurring during endocytic coat assembly *in vivo*, where it happens in the range of several seconds^1,2,38^.

**Fig. 6.**
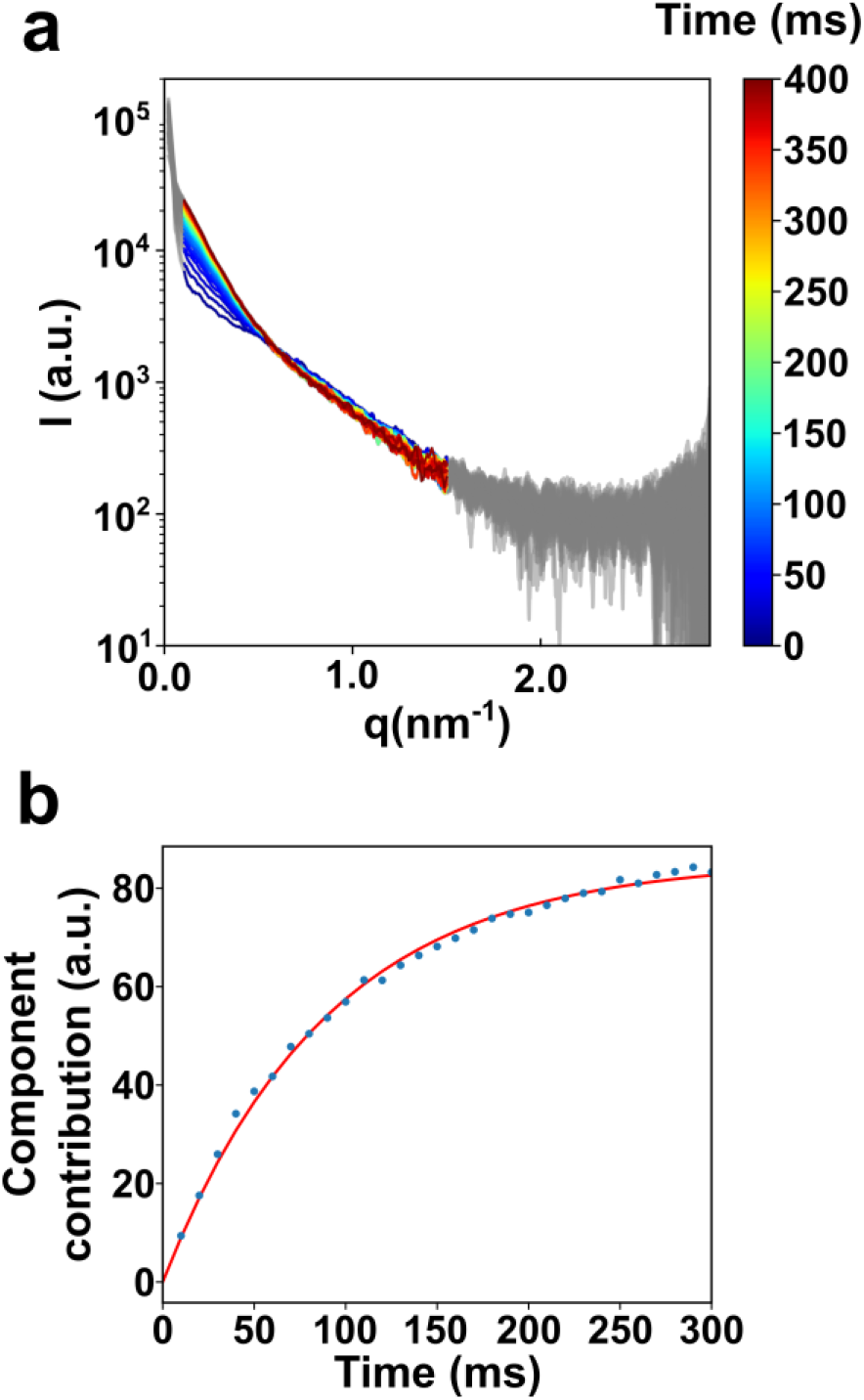
Time-resolved SAXS data demonstrate a fast assembly of ANTH and ENTH domains in solution. **a.** Buffer-corrected SAXS curves for the time-resolved measurements of the complex formation between ANTH and ENTH in presence of 200 μM PIP_2_. For better visibility, the data were smoothed with a Savitzky-Golay filter (51 point window, 2^nd^ order polynomial). Delay times are color coded from 0 ms in blue to 400 ms in dark red. The shape of the curve changes dramatically from the initial time points after 100 ms, when the curve stabilizes. **b.** The complex component contribution up to 300 ms was fitted to a single exponential curve showing a fast oligomerization of ANTH and ENTH domains in presence of PIP_2_.

Once endocytosis has been accomplished, the endocytic coat has to be disassembled and its components recycled. We performed biolayer interferometry (BLI) experiments, in which one of the adaptors was immobilized onto a surface and exposed to its partner in presence of PIP_2_, to obtain information regarding the reversibility of complex formation. Our results indicate that complex formation is fully reversible upon decreased concentrations of any of the two adaptor domains, since the signal decreases exponentially back to the baseline when the unattached adaptor domain was removed (**Table 3 and** Supplementary **Fig. 11**). This indicates that these domains are able to disassemble while the endocytic coat is being dismantled and the local concentration of one of their partner decreases.

**Table 3.** Fitting for the binding kinetics determined by BLI

| **Figure S11 A** |  |  |
| --- | --- | --- |
| **Fit rise** | **Fit decay** | **KD1 : 3.14E-08 M** |
| kobs1 : 3.06E-02 1/s | kdiss1 : 4.40E-03 1/s | **KD2 : 2.12E-07 M** |
| kobs2 : 3.45E-03 1/s | kdiss2 : 1.91E-02 1/s |  |
| R2 : 0.9989 | R2 : 0.9995 |  |

| **Figure S11 B** |  |  |
| --- | --- | --- |
| **Fit rise** | **Fit decay** | **KD1 : 1.93E-08M** |
| kobs1 : 7.45E-021s | kdiss1 : 6.23E-031s | **KD2 : 2.30E-07M** |
| kobs2 : 3.87E-031s | kdiss2 : 4.56E-021s |  |
| R2 : 0.9980 | R2 : 0.9981 |  |

## DISCUSSION

Clathrin-mediated endocytosis requires the concerted action of many proteins to coordinate cargo recruitment, endocytic coat formation and membrane bending. The early stages of the coat formation are essential for clathrin recruitment and proper assembly of the endocytic machinery for downstream actin polymerization and membrane invagination. We need to understand how the spatial and temporal regulation of CME functions to control relevant cellular processes involving adhesion complexes, signaling receptors, mechano-transducers, morphogen sensors, or polarity markers. Here, we determined the structure of a 16-mer assembly of the membrane-binding domains of Hip1R homolog Sla2 and epsin Ent1, ANTH and ENTH domains respectively, with PIP_2_ (8 ANTH:8 ENTH:20 PIP_2_) using cryo-EM. We found that the ANTH and ENTH domains arrange into tetrameric complexes that can further assemble into larger assemblies. The term di-octamer used in previous studies should therefore be re-interpreted as four tetramers^19,34^. The AENTH tetramer is also the building unit capable to remodel membranes during endocytosis, as shown for the ANTH-ENTH helical assembly on GUVs previously found^18^.

PIP_2_ was found in three different binding sites, one in the ENTH domain, previously known from the crystal structure^6^, one in the interface between the two ENTH domains of one tetramer, and one in the interface between the ANTH and ENTH domains, acting as a ‘glue’ for the oligomerization of both domains. The different binding sites present in the ENTH domain and especially shared binding sites between ANTH and ENTH domains agree with the positive cooperativity in the binding mechanism proposed before^18,19^.

The endocytic mechanism involves a plethora of protein-protein and protein-lipid interactions, however the strong affinity of these two domains for PIP_2_, which is specific to the endocytic site, makes it different from most other interactions in the endocytic coat. These are usually of low affinity to provide the coat with the dynamic characteristics required for this finely tuned biological process. The structure of A_8_E_8_ suggests that binding of the PIP_2_ to the ENTH domain, causing refolding of its α0 helix, is not sufficient to tightly anchor the ENTH domain and therefore epsin to the plasma membrane. A low affinity of ENTH to the membrane could be deleterious for endocytosis in yeast or in cells with high membrane tension, where CME depends on forces provided by actin and transmitted over the Sla2-epsin linker^21,22,39^. To achieve high affinity for PIP_2_-enriched membranes of endocytic sites, the ANTH domain must be present, sandwiching two PIP_2_ molecules shared with the ENTH domain and increasing thus the avidity of ANTH-ENTH tetrameric complex to the membrane. This tetrameric, PIP_2_-locked complex fits in the functional frame of these adaptors. It equips them to be anchored to the membrane strongly enough to transmit the force coming from the actin polymerization to the plasma membrane for membrane invagination to take place ^22^. Our structure shows the mechanism of lipid locking for this force transmission to take place, and how its efficiency relies sharing of multiple PIP_2_ molecules.

Our data suggest that ANTH and ENTH domains have evolved to achieve a fast assembly in presence of PIP_2_ and to do not require further proteins to form a stable complex. The whole endocytic process takes approximately 60 seconds^1^, and adaptors such as Sla2 and Ent1 assemble quickly and efficiently in the plasma membrane during the mid-coat formation to provide a scaffold for further assembly of the endocytic machinery. Therefore, it is biologically relevant that the kinetics of the oligomerization of these domains in solution show a very fast assembly of these essential pieces of the endocytic coat. Failure to do so would result in hampered endocytosis. An efficient disassembly should be achieved with the help of phosphatases, which at the end of the endocytic cycle remove PIP_2_ from the endocytic vesicle favouring the equilibrium of the oligomerization towards the disassembly of the endocytic adaptor complexes^1,24^. Our analysis of the different interfaces in the tetrameric assembly derived from the cryo-EM structure indicates that lack of impairment of some of these interfaces by introducing point mutations results in no complex formation and growth defect phenotypes *in vivo*.

In summary, ANTH and ENTH domains can assemble in a fast and coordinated way into their hetero-tetrameric functional unit, stabilized by shared PIP_2_ molecules. These tetramers can give rise to different assemblies depending on the lipid environment and membrane shapes occurring during endocytosis. The synergy of structural properties from ENTH α0 helices, combined with the lipid clamp of Sla2 ANTH domains makes the complex a better membrane anchor relying on shared PIP_2_ molecules in several protein interfaces. Thus, the combination of these two elements transforms the monomeric low-affinity (micromolar *Kd*) interaction with phospholipids^3,6^ into a nanomolar interaction^18,19^provided by a highly-organized protein-lipid-protein complex. This multimeric membrane anchor is then essential for membrane invagination in organisms and cells where endocytosis is challenged by high membrane tension or turgor pressure.

## MATERIALS AND METHODS

### Protein production and purification

Recombinant yeast Sla2 ANTH and Ent1 ENTH domains were expressed in *E. coli* BL21 DE3 (Novagen) as GST-fusion proteins containing an N-terminal His-tag followed by a TEV (Hisx6-TEV) cleavage site. Flasks containing 800 ml cultures in LB media were grown at 37 °C until an optical density (OD_600_) of 0.8. After induction with 0.5 mM IPTG, the cultures were grown at 20 °C for 4 h and harvested by centrifugation (4000 x g for 30 min at 4 °C). The cell pellet was lysed by sonication in the presence of 1 mg/ml DNase in 50 mM Tris-HCl pH 7.5, 300 mM NaCl, 20 mM imidazole. Lysed cell extract was centrifuged (17,000 x g, 45 min at 4 °C) and supernatant containing His-tagged proteins were purified by nickel-nitrilotriacetic acid (Ni-NTA) purification (Qiagen). Protein was eluted in a final elution buffer of 20 mM Tris pH 8.0, 300 mM NaCl, 250 mM imidazole. Excess of TEV protease was added to the imidazole-eluted fractions for cleavage of the His_6_-GST and His_6_ tags. Digestion was performed by dialysis at 4 °C overnight against 5 L of 20 mM Tris pH 8.0, 250 mM NaCl and 1 mM DTT. To remove the tags, the dialyzed fractions were subjected to a second Ni-NTA and the flow-through was concentrated to 5 mg/ml to be then injected in a size exclusion chromatography (SEC). SEC was performed using an ÄKTA liquid chromatography system (Amersham Biosciences) and S75 10/300 GL (Tricorn) column (GE Healthcare) in 20 mM Tris-HCl pH 8.0 and 250 mM NaCl. After SEC, the fractions were pooled and concentrated and flash frozen in liquid nitrogen and stored at −80 °C.

Lipid: di-C8-PIP2 was purchased from Echelon.

### Thermal denaturation assays

Proteins mixed to a final concentration of 30 μM in presence of 200 μM PIP_2_ and incubated overnight at 4 °C were used to fill two standard grade NanoDSF capillaries (Nanotemper) and loaded into a Prometheus NT.48 device (Nanotemper) controlled by PR.ThermControl (version 2.1.2). Excitation power was pre-adjusted to get fluorescence readings between 2000-20000 RFU for fluorescence at 330 and 350 nm (F330 and F350, respectively), and samples heated from 20 °C to 90 °C with a slope of 1 °C/min. An apparent *T_m_* was calculated from the inflection points of the fluorescence curves (Ratio F350/F330) for ANTH and ENTH monomeric samples, where Δ*Tm*=*Tm* mutant-*Tm* wild-type. Our results show that mutations of some conserved residues introduced protein destabilization effects: ANTH A15D and the double mutant ENTH F52A/F53A are clear examples of unstable domain mutants (Supplementary **Fig. 6**). To ensure we did not see any oligomerization interference effect potentially caused by domain instability, we discarded all mutants with a Δ*T_m_* larger than 2 °C in our complex assembly study.

The scattering signal recorded (back scattering mode) was used as a stability reporter for the AENTH samples, where the mid aggregation point, *T_agg_*, corresponds to the inflexion points in the scattering curves of the first transitions observed upon heating (T_agg_= mid-aggregation temperature obtained from scattering curves).

### Dynamic light scattering (DLS)

Measurements were performed using a DynaPro Nanostar device (Wyatt Technology Corporation) and data processed with Dynamics v.7 software. Proteins were mixed to a final concentration of 30 μM in presence of 200 μM PIP_2_ and incubated overnight at 4 °C. Samples were centrifuged at 10.000 rpm for 10 minutes prior to measurements using 4 μL plastic cuvettes (Wyatt Technology Corporation). The acquisition time was 5 s with a total of 30 acquisitions averaged. Measurements were performed at 25 °C.

### Stopped-flow time-resolved SAXS (SF-TR-SAXS) data collection and analysis

A stopped flow mixer (SFM 400, Bio-logic, Seyssinet-Pariset, France) equipped with a quartz capillary (0.8 mm inner diameter) was used as the sample delivery system at the SAXS beamline P12 (EMBL, PETRA III, DESY, Germany) ^41^. The device was used to rapidly mix ANTH and ENTH domains both in the presence of 200 μM PIP_2_ with a dead time of 5 ms.

SAXS curves were recorded with an EIGER X 4M detector at a distance of 3 m from the sample position using different time delays (0 ms, 400 ms, 800 ms, 1200 ms). These curves were collected in series spanning 600 ms and with a 200 ms overlap between each series. Each series contains spectra at 60 time points (10 ms spacing). Four data series with overlapping time points allowed to assess beam-induced effects on the sample. All spectra were solvent subtracted (buffer and PIP_2_ lipid) and used in the q-range between 0.01 and 2.9 nm^−1^ for further analysis. Singular value decomposition (SVD) of the processed spectra was performed by deconvolution of experimental data and kinetic fitting.

Based on the singular values, the number of species for the subsequent non-negative matrix factorization (NMF) was chosen to be three. The NMF component spectra were assigned to complex, monomers and radiation damage. The component contributions of the complex up to 300 ms time delay were fit to a single exponential. For all steps, self-written Python code using the modules Numpy ^42^, Scipy ^43^ and Scikit-learn ^44^ was used.

### Grid preparation

ANTH and ENTH at a concentration of 100 μM were pre-incubated in 200 μM PIP_2_ in buffer containing 20mM Tris pH 8.0, 250 mM NaCl and 1 mM DTT for 3 h at room temperature. Then, the solutions were mixed 1:1 to generate the AENTH complex and left on ice for at least 1 h. For cryo-EM grid preparation, Quantifoil 300 mesh Cu R 1.2/1.3 holey carbon grids were glow-discharged in a Cressington 208 carbon coater at 10 mA and 0.1 mbar air pressure for 30 s. The complex was diluted to 10 μM ANTH/ENTH (monomer) in 200 μM PIP_2_ and 3 μL was then applied to the grid and vitrified using a Vitrobot™ mark IV (Thermo/FEI) with a blot force of 6 and a blot time of 6 s. The relative humidity (RH) was ≥ 90% and temperature 5-6 °C. Liquid ethane was used as the cryogen.

### Data collection and processing

Cryo-EM data were collected on a Titan Krios (Thermo/FEI) at the Astbury Biostructure Laboratory using a Falcon III direct electron detector operating in integrating mode. The main data acquisition parameters for the wildtype dataset (A_8_E_8_) are listed in Table 1. Processing of the (A_2_E_2_)_4_ data was done using RELION 3^45^ and cryoSPARC v2: Micrographs were corrected for beam induced motion using MotionCor2^46^ and the contrast transfer function (CTF) was estimated using Gctf^47^, in RELION. Particles were picked initially using the general model in crYOLO^48^ to generate initial 2D classes and a 3D reconstruction in RELION. The model was then trained to pick 16-mer particles. Using the trained model, 195,536 particles were picked from 7,990 micrographs. A subset of these was used to generate an initial model and all particles were subjected to 3D classification to remove ‘bad’ particles. 96,664 particles were taken forward to refinement in C1 and D2 symmetry. Bayesian polishing^49^ and beamtilt estimation were applied and a 2D classification step was performed on the polished particles to give a final selection of 79,414 particles, leading to a resolution of 4.1 Å (D2 symmetry). Non-uniform refinement in cryoSPARC v2^50^ was used to further improve the resolution to 3.9 Å and the final reconstruction was sharpened with B-factor of −200. An overview of the processing strategy is given in Supplementary **Fig. 2**.

Data collection parameters for the F5A/L12A/V13A mutant (A_6_E_6_) assembly are listed in Supplementary **Table 1**. Processing of the (A_2_E_2_)_3_ data was done in RELION3 (Zivanov et al., 2018). Motion-correction and CTF-estimation were as for the (A_2_E_2_)_4_ data. Using the general model in crYOLO, 142,399 particles were picked from 1,791 micrographs. Those were subjected to two rounds of 2D-classification, and a subset of 16,206 particles was selected. An initial model was generated from these particles, imposing C2 symmetry. The model was then refined with D3 symmetry, and particles were subjected to Bayesian polishing, CTF refinement and beamtilt estimation, yielding a final reconstruction with 7.4 Å global resolution. An overview of the processing strategy is given in Supplementary Fig. 10.

### Model building

As initial model for the *S. cerevisiae* Sla2 ANTH domain, the *Chaetomium thermophilum* Sla2 ANTH domain crystal structure (PDB: 5oo7) was used. This, and the *S. cerevisiae* ENTH domain crystal structure (PDB: 5onf) were rigid-body docked into the cryo-EM map of A_8_E_8_ using Chimera and manually adjusted in Coot ^51^. The tetramer occupying the asymmetric unit was then iteratively refined in Coot and ISOLDE ^52^. In total, 20 PIP_2_ ligands were identified and placed in the model in Coot. Ligand coordinates and restraints were generated, symmetry applied and validation performed using tools in PHENIX 1.17 ^53^.

For comparison with the previously published structure of ANTH-ENTH on lipid tubules, the atomic model for the A_2_E_2_ tetramer was docked into a subvolume of the EMD-2896 map and then flexibly fitted using adaptive distance restraints in ISOLDE.

Cryo-EM maps and atomic models were visualized using Chimera and ChimeraX^54,55^.

### Native mass spectrometry

Borosilicate nano-electrospray capillaries (Thermo Scientific) were prepared in-house using a P-97 micropipette puller (Sutter Instrument Co.), and coated with palladium/gold in a Polaron SC7620 sputter coater (Quorum Technologies). Native mass spectrometry measurements were done using an Orbitrap Q Exactive Plus UHMR (Thermo Scientific) operated in positive ion mode. Proteins were buffer-exchanged using one or two consecutive Zeba spin desalting columns (Thermo Scientific) into 300 mM ammonium acetate, 1 mM DTT, pH 8. Then, the desired amount of PIP_2_ was added to a mixture of ANTH and ENTH proteins (1:1 molar ratio) to form the complex giving a final (monomer) concentration of 10 μM. Instrument settings were 1.5-1.6 kV capillary voltage, −150 V in source trapping, HCD was off and the AGC target set to 3×10^6^ with a maximum inject time of 300 ms. The trapping gas pressure (ratio) was 8, the mass range was 2000 – 20000 m/z and the resolution set to 6,250. Raw data were processed and analyzed using UniDec^56^.

### Biolayer interferometry

Measurements were performed using an Octet RED96 instrument and Ni-NTA biosensors (ForteBio). Protein solutions were centrifuged for 10 min at 10,000 rpm at 4°C before the experiment to remove possible aggregates. Protein concentrations of these stock solutions were determined after centrifugation by the absorbance at 280 nm with a NanoDrop1000. Prior to the experiment, the biosensors were equilibrated in buffer I (50 mM Tris, pH 8; 125 mM NaCl and 0.05% BSA) for 10 min. Binding between His-T_agg_ed ANTH and ENTH from *Chaetomium thermophilum* was measured in the presence of 170 *μ*M DDM and 50 *μ*M PIP_2_ in buffer I (buffer II). Prior attempts to use PIP_2_ (225*μ*M) without DDM resulted in unspecific binding of ENTH to the biosensor. The experiments were performed at 25°C with a shaking speed of 1000 rpm. An evaporation cover was used throughout the experiment.

Kinetic assays were performed in black, flat-bottom polypropylene 96 well plates (Greiner bio-one, item no. 655201) using 200 *μ*l in each well. The kinetic assay consisted of 4 steps: 300 s equilibration in buffer I (baseline I), 300s loading (3.75 *μ*g/ml His-tagged ANTH in buffer I), 300 s equilibration in buffer II (baseline II), 600 s ENTH association (0.25 *μ*M ENTH in buffer II), 1200 s dissociation in buffer II. To test for unspecific binding, a kinetic assay without loading His-tagged ANTH was used as control.

Data were visualized and analyzed with self-written Python scripts using the Python packages Numpy ^42,57^, Matplotlib ^58^ and Scipy ^43^. Association and dissociation were fit with a biexponential function yielding a pair of two *kobs* and *kdiss*, each.

## Acknowledgements

We would like to acknowledge the Sample Preparation and Characterization Facility at EMBL Hamburg. Synchrotron SAXS data was collected at beamline P12 operated by EMBL Hamburg at the PETRA III storage ring (DESY, Hamburg, Germany). We would like to thank beam scientists for the assistance in using the beamline. The work of M.A. and M.S. was supported by Deutsche Forschungsgemeinschaft (DFG) Research Grant SK 305/1-1. D.P.K. is a PhD student on the Wellcome Trust 4-year PhD programme in The Astbury Centre funded by The University of Leeds. The FEI Titan Krios microscope was funded by the University of Leeds (UoL ABSL award) and Wellcome Trust (108466/Z/15/Z). The Orbitrap UHMR was funded by the University of Leeds and Wellcome Trust multi-user equipment grant 208385/Z/17/Z.

## Author contributions

J.L. and K.V. produced proteins and samples for cryo-EM, SAXS, and native Mass spectrometry. J.L. performed biophysical experiments and interpreted EM Data. D.K. performed cryo-EM experiments and determined the AENTH structures together with S.M.. D.K. performed mass spectrometry experiments and interpreted the data together with F.S. M.A. and M.S. performed growth experiments in S. *cerevisiae* and interpreted data. M.A.S. and H.M. performed SAXS experiments and interpreted the data together with D.I.S.. S.N. performed BLI experiments and SF-TR-SAXS data analysis with input from M.A.S.. M.G.A. conceived and supervised the project. J.L. and M.G.A. wrote the manuscript with input from all authors.

## Additional information

**Supplementary Information** is available for this paper. Correspondence and requests for materials should be addressed to M.G.A.

## Supplementary Figures

**Supplementary Fig. 1:**
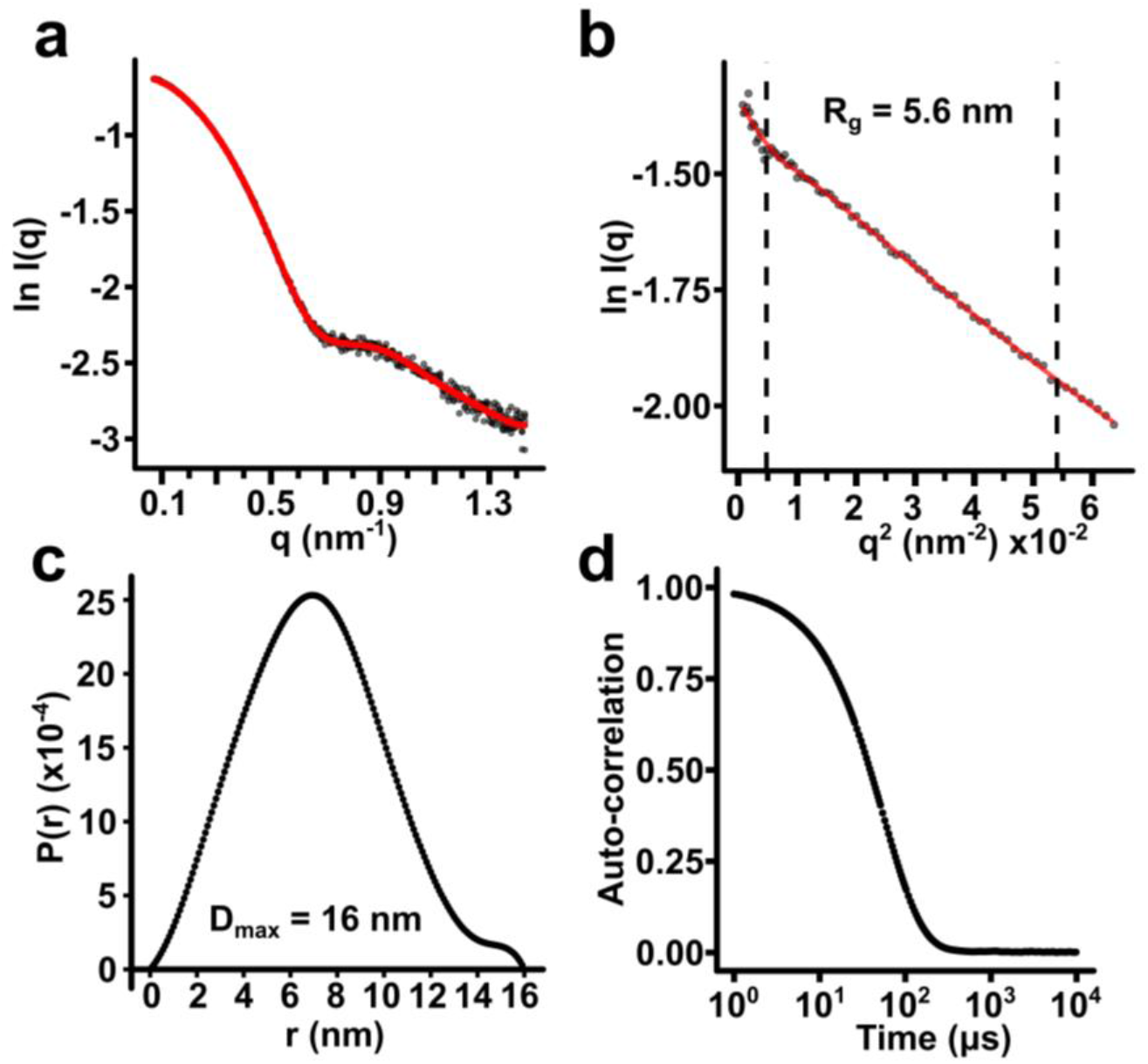
Biophysical characterization of the AENTH assembly in solution. **a.** Experimental SAXS data of the 16-mer assembly (A_8_E_8_) in presence of 200 μM PIP_2_. The red line represents the fitting of a mixture population in which the major species is the A_8_E_8_ assembly. **b.** Guinier plot for calculation of the R_g_ of the 16-mer assembly. **c.** Distance-distribution function for the 16-mer assembly. **d.** DLS auto-correlation curve for the sample used for cryoEM.

**Supplementary Fig. 2:**
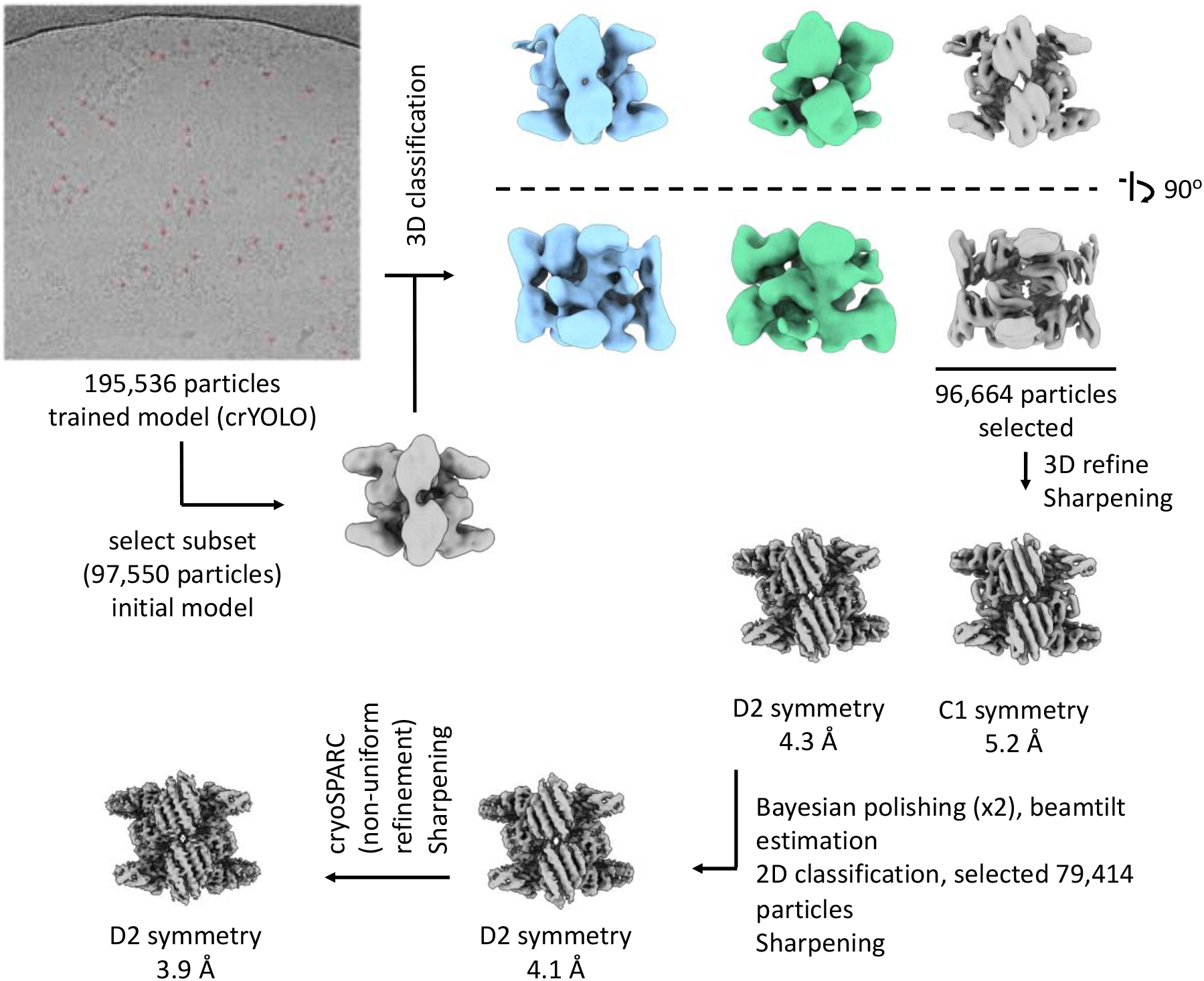
Processing flowchart for the 16-mer AENTH complex.

**Supplementary Fig. 3.**
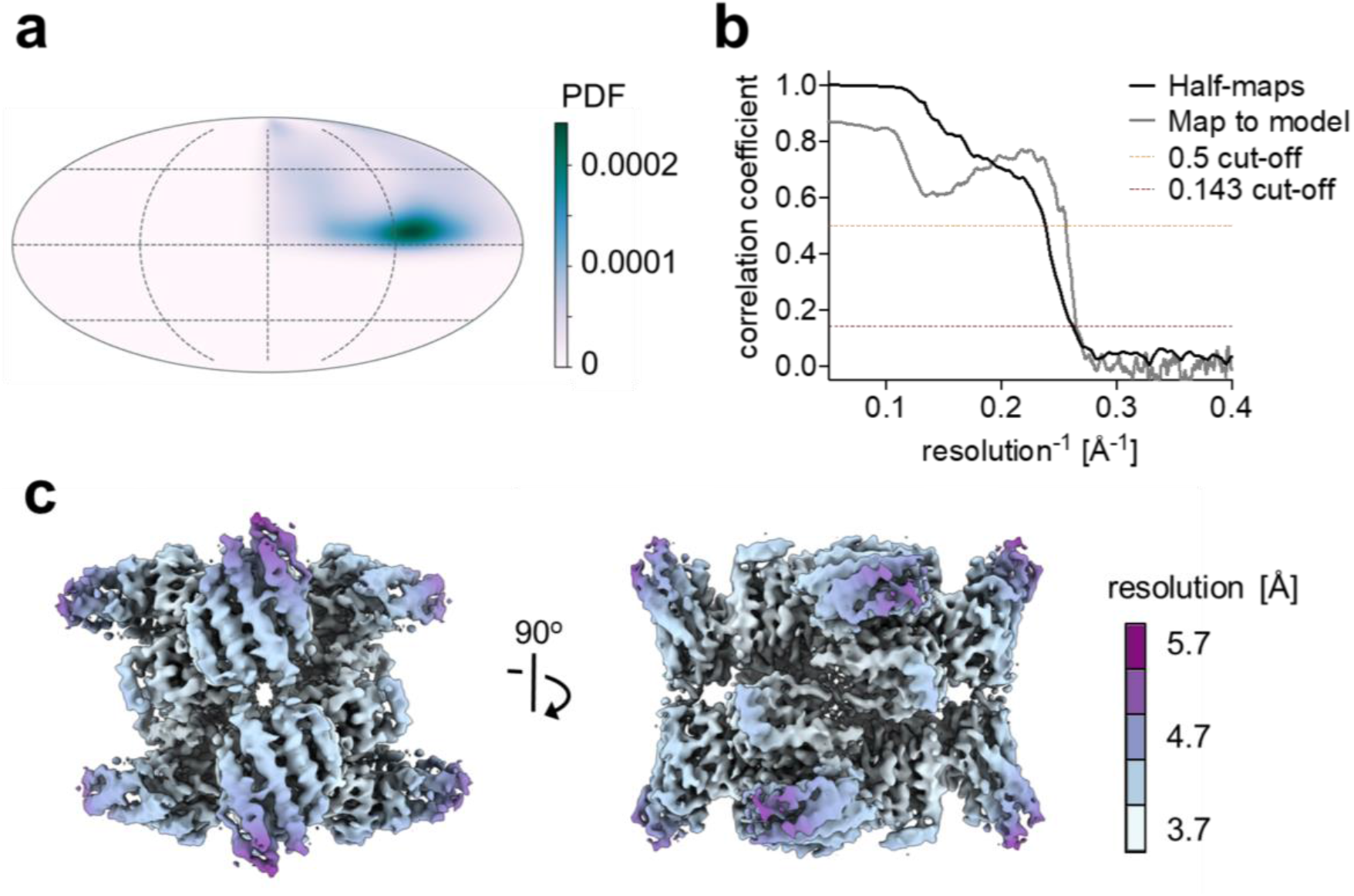
**a.** Angular distribution. **b.** Fourier Shell Correlation (FSC) curves and **c.** local resolution for the AENTH 16-mer complex.

**Supplementary Fig. 4.**
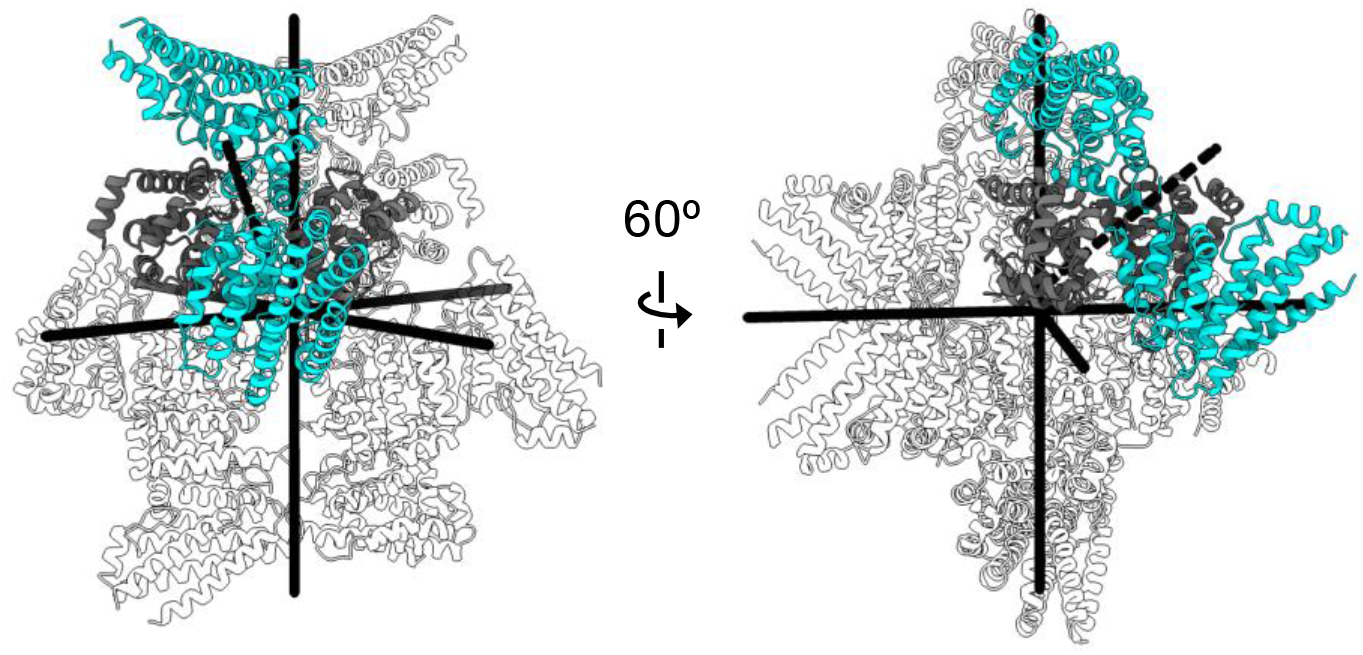
Structure of 16-mer AENTH (A_8_E_8_) complex showing symmetry axis. Now also showing The pseudo-2fold axis within the tetramer is shown as dashed line.

**Supplementary Fig. 5.**
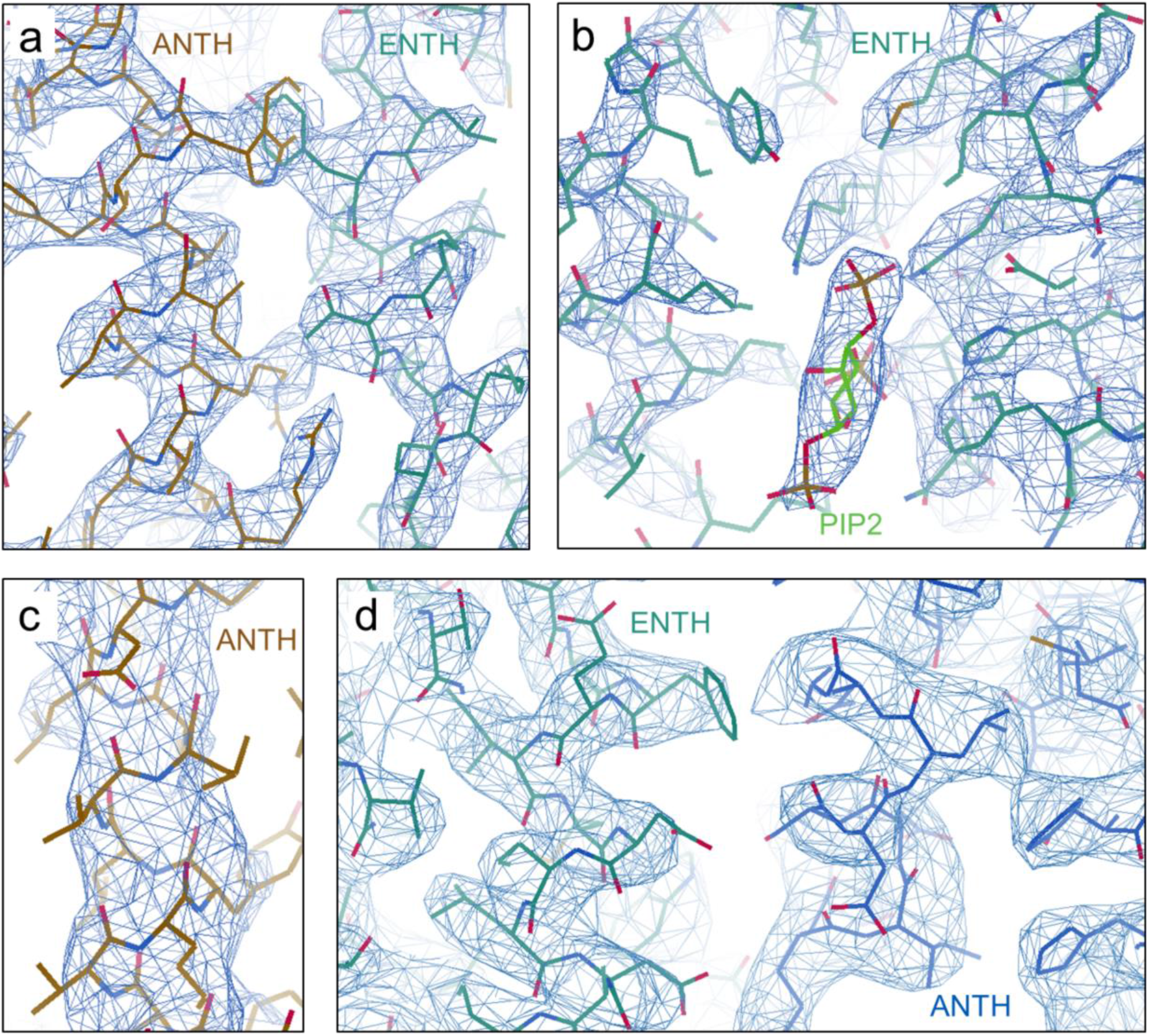
Representative EM density of (**a**) well-resolved central alpha-helical parts of the ANTH-ENTH complex, (**b**) one of the PIP_2_-binding sites in between two ENTH domains, (**c**) lower resolution at the periphery of the complex in the last alpha helix of the ANTH domain and (**d**) in the ANTH-ENTH interface 2.

**Supplementary Fig. 6.**
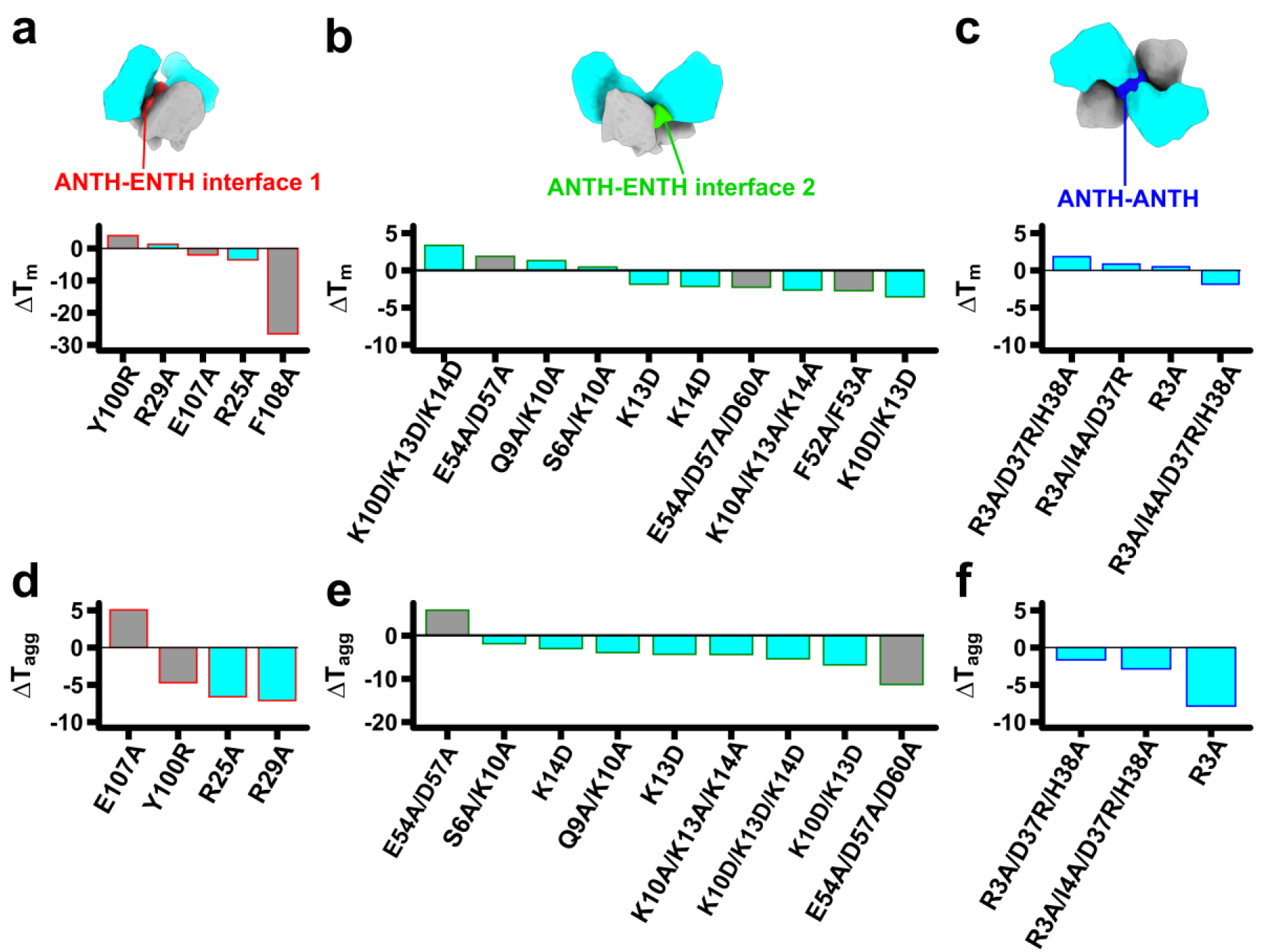
nanoDSF data for the ANTH and ENTH mutants affecting ANTH-ENTH interface 1 (red), ANTH-ENTH interface 2 (green) and ANTH-ANTH interface (blue) (**a, b and c)**: Difference in melting temperature (T_m_) between mutant domains compared to wild-type ANTH (cyan) or ENTH (grey). The T_m_ was obtained from the fluorescence signal ratio 350/330 nm. All domains with a ΔT_m_ higher than 2 °C were not further considered for complex assembly experiments. (**d, e and f)**: Difference in the mid-aggregation temperature (T_agg_) from AENTH complexes assembled with different mutant domains (ANTH, cyan and ENTH, grey) compared to wild-type AENTH. T_agg_ was obtained from the scattering signal of the nanoDSF melting experiments. Most mutants showed a lower aggregation temperature when compared with wild-type.

**Supplementary Fig. S7.**
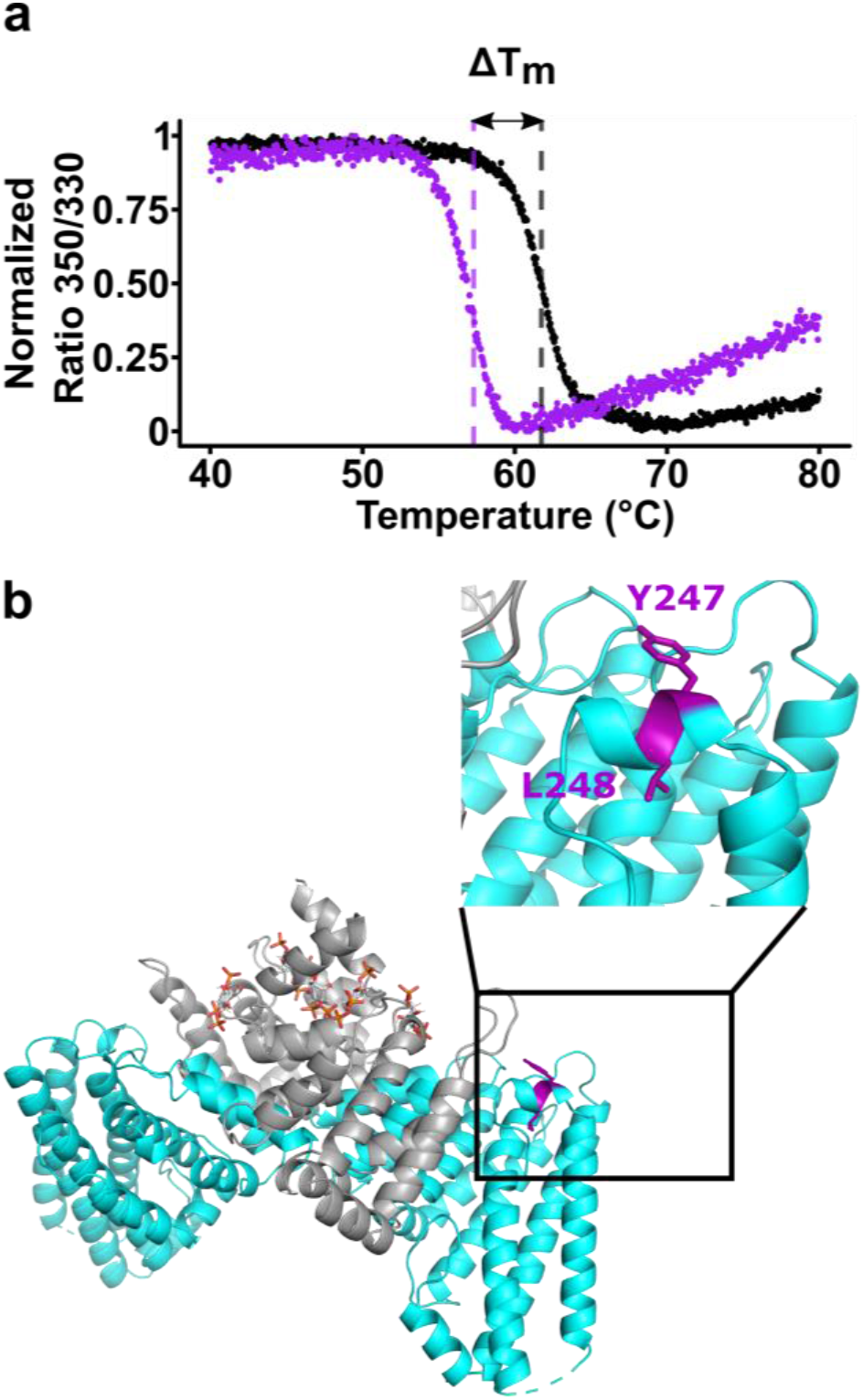
**a.** nanoDSF data for ANTH WT (black, T_m_= 61.7 °C) and ANTH ΔY247/L248 (violet, T_m_ = 57.7 °C). The shift in the transition of the nanoDSF signal can be clearly observed and is indicated as ΔT_m_, indicating that this mutation destabilizes the protein. The T_m_ was obtained from the fluorescence signal ratio 350/330 nm. **b.** Residues Y247 and L248 of ANTH domain did not establish any contacts in the AENTH complex. Cartoon representation of the AENTH tetramer and the PIP_2_ molecules bound to it, with the YL residues colored in violet. The insert shows the detailed orientation of these residues, which doesn’t establish any protein-protein contacts. Deletion of these residues disrupts helix α12.

**Supplementary Fig. 8.**
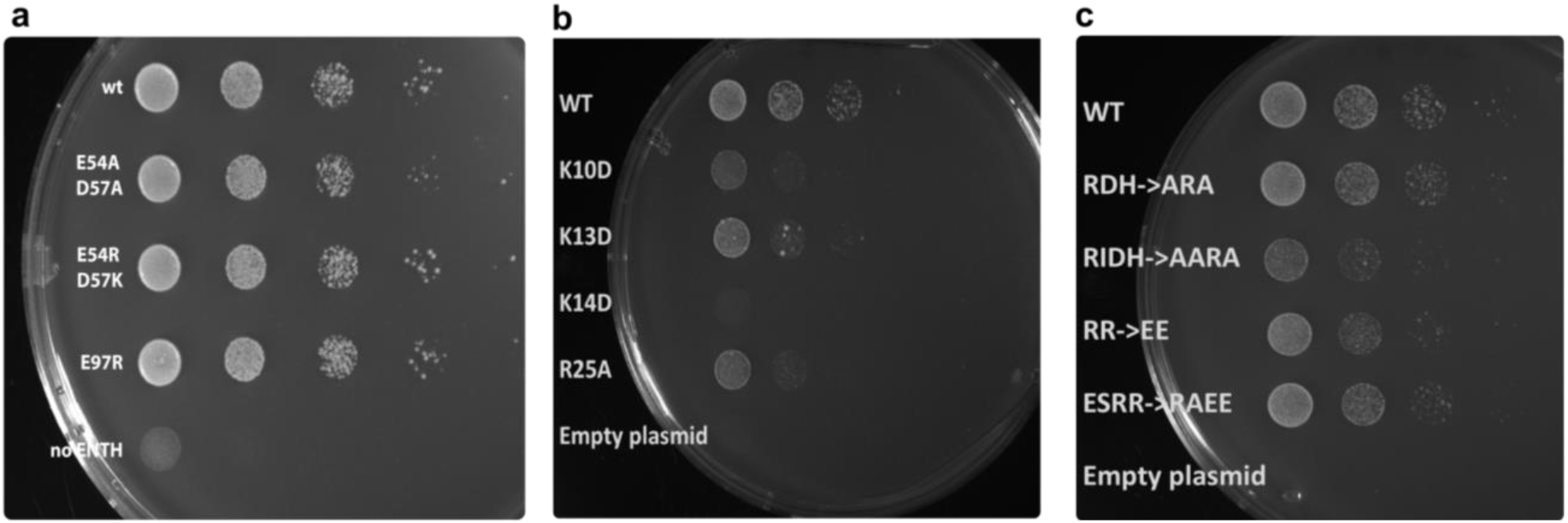
*In vivo* results for selected mutations affecting the ANTH-ENTH interface 2 (**a** and **b**) and the ANTH-ANTH interface (**c**). Cell growth was analyzed by plating 10-fold serial dilution of cells on SD-Ura plates and incubated for 3 days at 37 °C. **a.** Mutation of negatively charged residues on ENTH E54 and D57 does not introduce growth defects. **b.** Mutation of single ANTH positively charged residues K10, K13, K14 and R25 impair efficient cell growth. **c.** Effect of mutants on the ANTH-ANTH interface (RDH = R3A/D37R/H38A, RIDH = R3A/I4A/D37R/H38A) and the ANTH loop 175-183 (RR = R178E/E178E, ESRR = E57R/S100A/R178E/E178E).

**Supplementary Fig. 9.**
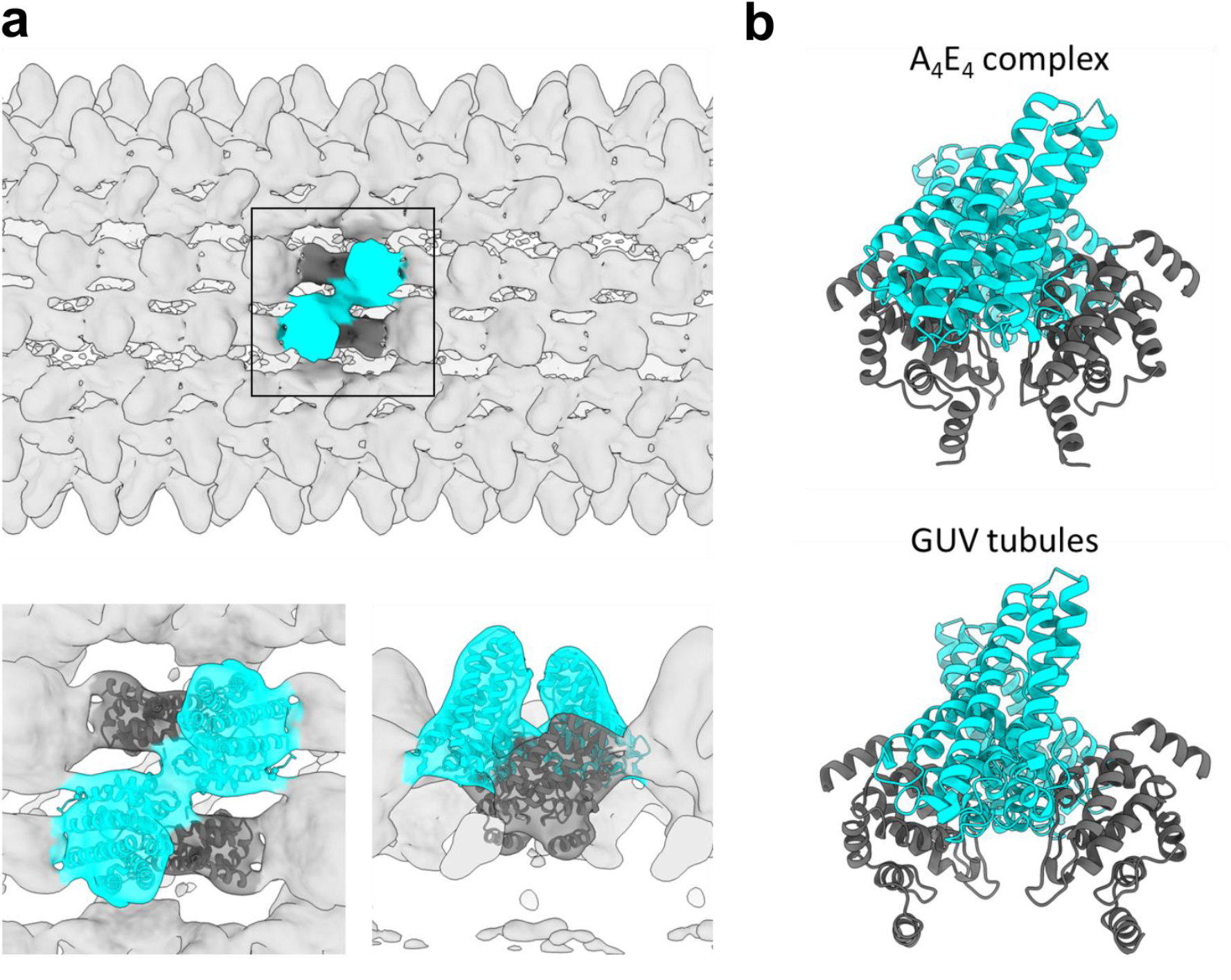
Flexible fitting of the tetrameric (A_2_E_2_) model to the previously obtained cryoEM map of ANTH-ENTH coat on GUV tubules (Skruzny et al., 2015). **a. (top)** Overview of the GUVs tubular coat structure with the tetramer fitting inside the lobes of the structure (EM-DB entry: EM-2896). **(bottom)** Close-up view of the tetramer fitted into the EM density of the GUV structure. In agreement with previous observations, the ANTH domains correspond to the larger density while the ENTH is in the interior of the tubular structure. Also, the α0 helix is pointing towards the interior of the structure. **b.** Comparison of the tetramer in the 16-mer ANTH-ENTH complex and in the fitted structure to the GUVs coat structure.

**Supplementary Fig. 10.**
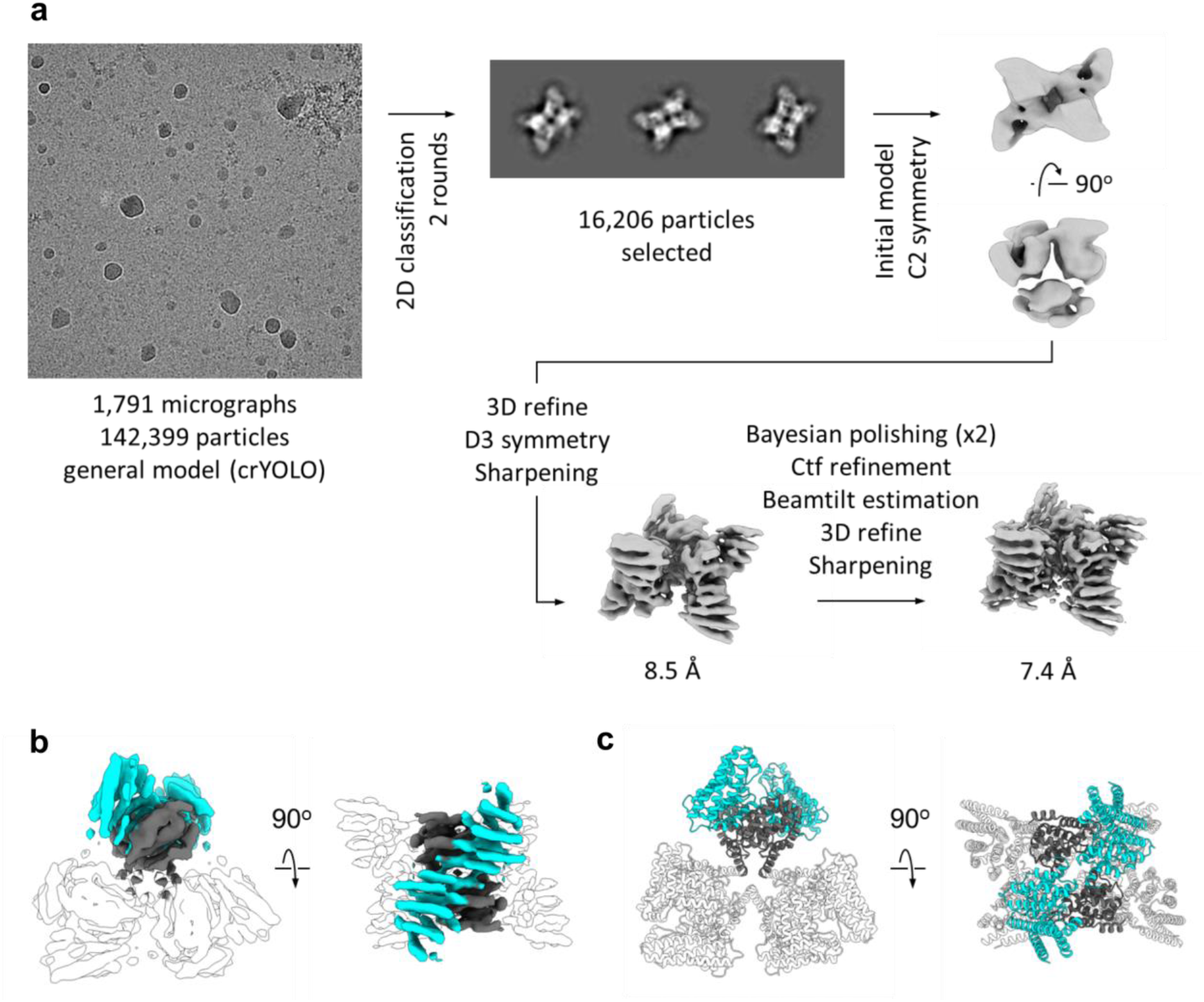
AENTH 12-mer assembly (A_6_E_6_) obtained by mutation of residues F5A/L12A/V13A of the amphipatic α0 helix of the ENTH domain. **a.** Processing flowchart for the 12-mer AENTH complex. **b**. Final density map obtained for the 12-mer assembly with the different subunits coloured in cyan for the ANTH domain and in grey for the ENTH domain. **c**. Structural model for the 12-mer assembly. The structure from the A_2_E_2_ was fitted into the EM density map for each of the three tetramers. The ANTH and ENTH domains are coloured for one of the three tetramers in the same colour code as in **b**.

**Supplementary Fig. 11.**
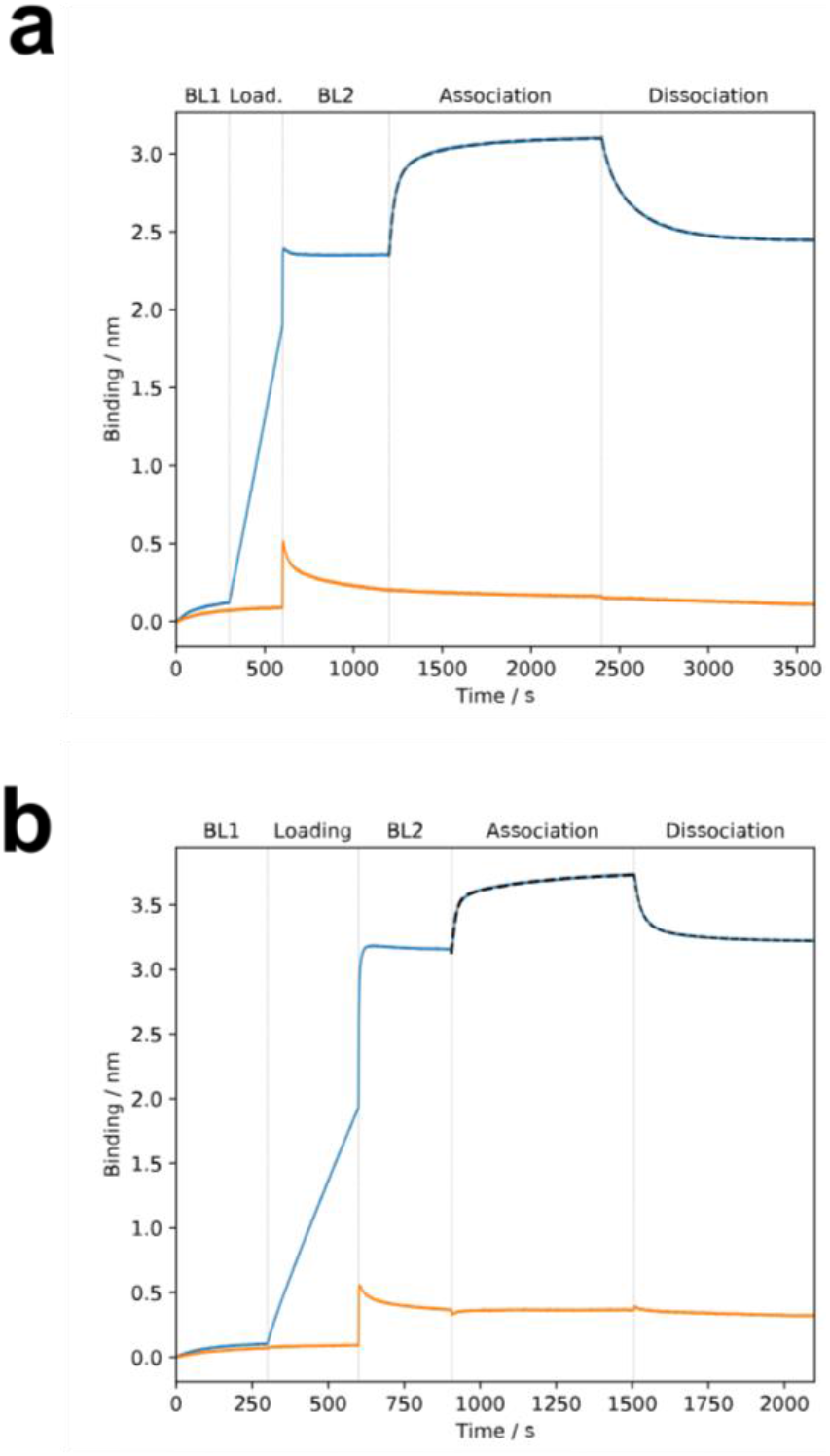
AENTH complex formation is a reversible process. Binding kinetics measured by biolayer interferometry (BLI) between His-tagged ANTH (**a**) or ENTH (**b**) immobilized on a Ni-NTA Octet sensor (load stage performed at 3.7 μg/ml monomeric protein) in the presence of 0.25 μM free ENTH (**a**) or ANTH (**b**) (blue curves) and without ligand (orange curves). All steps were done in 50 mM Tris HCl pH 8.0, 125 mM NaCl and 0.05% BSA. For baseline 2 (BL2), association and dissociation stages the buffer additionally contained 170 μM DDM and 50 μM PIP_2_. See Table 5 for the kinetic constants.

**Supplementary Table 1:** Data collection and processing parameters for the ANTH-ENTH A_6_E_6_ complex.

| **Data collection and processing** |  |
| --- | --- |
| Magnification | ×75,000 |
| Voltage (kV) | 300 |
| Electron exposure (e<sup>-</sup>/Å<sup>2</sup>) | 72.6 |
| Defocus range (μm) | -1.5 to -4.2 |
| Pixel size (Å) | 1.065 |
| Symmetry imposed | D<sub>3</sub> |
| Initial particle images (no.) | 142,399 |
| Final particle images (no.) | 16,206 |
| Map resolution (Å) | 7.4 |
| FSC threshold | 0.143 |
| Map resolution range (Å) | 6.8 to 9.9 |

## REFERENCES

1. Kaksonen, M. & Roux, A. Mechanisms of clathrin-mediated endocytosis. Nat. Rev. Mol. Cell Biol. (2018). doi:10.1038/nrm.2017.132

2. Lu, R., Drubin, D. G. & Sun, Y. Clathrin-mediated endocytosis in budding yeast at a glance. J. Cell Sci. 129, 1531–1536 (2016).

3. Smith, S. M., Baker, M., Halebian, M. & Smith, C. J. Weak Molecular Interactions in Clathrin-Mediated Endocytosis. 4, (2017).

4. Edeling, M. A. et al. Molecular switches involving the AP-2 β2 appendage regulate endocytic cargo selection and clathrin coat assembly. Dev. Cell 10, 329–342 (2006).

5. Ford, M. G. J. et al. Simultaneous binding of PtdIns (4,5) P 2 and clathrin by AP180 in the nucleation of clathrin lattices on membranes. Science (80-.). 291, 1051–1055 (2001).

6. Ford, M. G. J. et al. Curvature of clathrin-coated pits driven by epsin. Nature 419, 361–366 (2002).

7. Kalthoff, C., Alves, J., Urbanke, C., Knorr, R. & Ungewickell, E. J. Unusual structural organization of the endocytic proteins AP180 and epsin 1. J. Biol. Chem. 277, 8209–8216 (2002).

8. Miller, S. E. et al. CALM Regulates Clathrin-Coated Vesicle Size and Maturation by Directly Sensing and Driving Membrane Curvature. Dev. Cell 33, 163–175 (2015).

9. Bucher, D. et al. Clathrin-Adaptor ratio and membrane tension regulate the flat-To-curved transition of the clathrin coat during endocytosis. Nat. Commun. 9, (2018).

10. Kukulski, W., Schorb, M., Kaksonen, M. & Briggs, J. A. G. Plasma membrane reshaping during endocytosis is revealed by time-resolved electron tomography. Cell 150, 508–520 (2012).

11. Morris, K. L. et al. universal mode of clathrin self-assembly. Nat. Struct. Mol. Biol. 26, (2019).

12. Scott, B. L. et al. Membrane bending occurs at all stages of clathrincoat assembly and defines endocytic dynamics. Nat. Commun. 9, (2018).

13. Kaksonen, C. J. M. and M. Endocytic Accessory Factors and Regulation Clathrin-Mediated Endocytosis. 1–16 (2018).

14. Mund, M. et al. Systematic Nanoscale Analysis of Endocytosis Links Efficient Vesicle Formation to Patterned Actin Nucleation. Ssrn 884–896 (2018). doi:10.2139/ssrn.3155759

15. Picco, A. & Kaksonen, M. Precise tracking of the dynamics of multiple proteins in endocytic events. Methods in Cell Biology 139, (Elsevier Ltd, 2017).

16. Payne, G. S., Baker, D., Evert van Tuinen & Schkman, R. Protein Transport to the Vacuole and Receptor-mediated Endocytosis by Clathrin Heavy Chain-deficient Yeast. 106, 1453–1461 (1988).

17. Kukulski, W., Picco, A., Specht, T., Briggs, J. A. G. & Kaksonen, M. Clathrin modulates vesicle scission, but not invagination shape, in yeast endocytosis. Elife 5, 1–10 (2016).

18. Skruzny, M. et al. An Organized Co-assembly of Clathrin Adaptors Is Essential for Endocytosis. Dev. Cell 33, 150–162 (2015).

19. Garcia-Alai, M. M. et al. Epsin and Sla2 form assemblies through phospholipid interfaces. Nat. Commun. 9, 1–13 (2018).

20. Engqvist-Goldstein, Å. E. Y. et al. The actin-binding protein Hip1R associates with clathrin during early stages of endocytosis and promotes clathrin assembly in vitro. J. Cell Biol. 154, 1209–1223 (2001).

21. Messa, M. et al. Epsin deficiency impairs endocytosis by stalling the actin-dependent invagination of endocytic clathrin-coated pits. Elife 3, 1–25 (2014).

22. Skruzny, M. et al. Molecular basis for coupling the plasma membrane to the actin cytoskeleton during clathrin-mediated endocytosis. Proc. Natl. Acad. Sci. 109, E2533–E2542 (2012).

23. Yang, S., Cope, M. J. T. V & Drubin, D. G. Sla2p Is Associated with the Yeast Cortical Actin Cytoskeleton via Redundant Localization Signals. 10, 2265–2283 (1999).

24. Di Paolo, G. & De Camilli, P. Phosphoinositides in cell regulation and membrane dynamics. Nature 443, 651–657 (2006).

25. Sun, Y. & Drubin, D. G. The functions of anionic phospholipids during clathrin-mediated endocytosis site initiation and vesicle formation. J. Cell Sci. 125, 6157–6165 (2012).

26. Campelo, F., Mcmahon, H. T. & Kozlov, M. M. The Hydrophobic Insertion Mechanism of Membrane Curvature Generation by Proteins. 95, 2325–2339 (2008).

27. Haucke, V. & Kozlov, M. M. Membrane remodeling in clathrin-mediated endocytosis. J. Cell Sci. 131, 1–10 (2018).

28. Kozlov, M. M. et al. Mechanisms shaping cell membranes. Curr. Opin. Cell Biol. 29, 53–60 (2014).

29. Yoon, Y. et al. Molecular basis of the potent membrane-remodeling activity of the epsin 1 N-terminal homology domain. J. Biol. Chem. 285, 531–540 (2010).

30. Kweon, D. H. et al. Membrane topology of helix 0 of the epsin N-terminal homology domain. Mol. Cells 21, 428–435 (2006).

31. Lai, C. et al. Membrane Binding and Self-Association of the Epsin N-Terminal Homology Domain. J. Mol. Biol. 423, 800–817 (2012).

32. Itoh, T. et al. Role of the ENTH domain in phosphatidylinositol-4,5-bisphosphate binding and endocytosis. Science (80-.). 291, 1047–1051 (2001).

33. Aguilar, R. C. et al. Epsin N-terminal homology domains perform an essential function regulating Cdc42 through binding Cdc42 GTPase-activating proteins. Proc. Natl. Acad. Sci. U. S. A. 103, 4116–4121 (2006).

34. Heidemann, J. et al. Further insights from structural mass spectrometry into endocytosis adaptor protein assemblies. Int. J. Mass Spectrom. 447, 1–9 (2020).

35. McMahon, H. T. & Boucrot, E. Membrane curvature at a glance. J. Cell Sci. 128, 1065–1070 (2015).

36. Walsh, J. P., Suen, R., Glomset, J. A. & Chem, B. Specific in vitro inihibition by phosphoinositides suggests a mechanism for regulation of phosphatidylinositol biosinthesis. 270, 28647–28653 (1995).

37. Weigang Huang, Dechen Jiang, Xiaoyang Wang, Kelong Wang, Christopher E. Sims, N. L. A. and Q. Z. NIH Public Access. 401, 1881–1888 (2012).

38. Brach, T., Godlee, C., Moeller-hansen, I., Boeke, D. & Kaksonen, M. Report The Initiation of Clathrin-Mediated Endocytosis Is Mechanistically Highly Flexible. Curr. Biol. 24, 548–554 (2014).

39. Boulant, S., Kural, C., Zeeh, J. C., Ubelmann, F. & Kirchhausen, T. Actin dynamics counteract membrane tension during clathrin-mediated endocytosis. Nat. Cell Biol. 13, 1124–1132 (2011).

40. Taneva, S. G., Lee, J. M. C. & Cornell, R. B. The amphipathic helix of an enzyme that regulates phosphatidylcholine synthesis remodels membranes into highly curved nanotubules. Biochim. Biophys. Acta - Biomembr. 1818, 1173–1186 (2012).

## Methods References

1. Josts, I. et al. Structural Kinetics of MsbA Investigated by Stopped-Flow Time-Resolved Small-Angle X-Ray Scattering. Structure 28, 348–354.e3 (2020).

2. Oliphant, T. E. Guide to NumPy. Methods 1, 378 (2010).

3. Virtanen, P. et al. Author Correction: SciPy 1.0: fundamental algorithms for scientific computing in Python (Nature Methods, (2020), 17, 3, (261-272), 10.1038/s41592-019-0686-2). Nat. Methods 17, 352 (2020).

4. Pedregosa, F. et al. Scikit-learn: Machine learning in Python. J. Mach. Learn. Res. 12, 2825–2830 (2011).

5. Zivanov, J. et al. New tools for automated high-resolution cryo-EM structure determination in RELION-3. Elife 7, 1–22 (2018).

6. Zheng, S. Q. et al. MotionCor2 : anisotropic correction of beam-induced motion for improved cryo-electron microscopy Automatic tracing of ultra-volumes of neuronal images. 14, 331–332 (2017).

7. Zhang, K. Gctf: Real-time CTF determination and correction. J. Struct. Biol. 193, 1–12 (2016).

8. Wagner, T. et al. SPHIRE-crYOLO is a fast and accurate fully automated particle picker for cryo-EM. Commun. Biol. 1–13 doi:10.1038/s42003-019-0437-z

9. Zivanov, J., Nakane, T. & Scheres, S. H. W. A Bayesian approach to beam-induced motion correction in cryo-EM single-particle analysis. 5–17 (2019). doi:10.1107/S205225251801463X

10. Punjani, A., Zhang, H. & Fleet, D. J. Non-uniform refinement: Adaptive regularization improves single particle cryo-EM reconstruction. bioRxiv 2019.12.15.877092 (2019). doi:10.1101/2019.12.15.877092

11. Emsley, P., Lohkamp, B., Scott, W. G. & Cowtan, K. Features and development of Coot. Acta Crystallogr. Sect. D Biol. Crystallogr. 66, 486–501 (2010).

12. Croll, T. I. ISOLDE: A physically realistic environment for model building into low-resolution electron-density maps. Acta Crystallogr. Sect. D Struct. Biol. 74, 519–530 (2018).

13. Liebschner, D. et al. Macromolecular structure determination using X-rays, neutrons and electrons: Recent developments in Phenix. Acta Crystallogr. Sect. D Struct. Biol. 75, 861–877 (2019).

14. Pettersen, E. F. et al. UCSF Chimera - A visualization system for exploratory research and analysis. J. Comput. Chem. 25, 1605–1612 (2004).

15. Goddard, T. D. et al. UCSF ChimeraX: Meeting modern challenges in visualization and analysis. Protein Sci. 27, 14–25 (2018).

16. Marty, M. T. et al. Bayesian deconvolution of mass and ion mobility spectra: From binary interactions to polydisperse ensembles. Anal. Chem. 87, 4370–4376 (2015).

17. van der Walt, S. J., Colbert, S. C. & Varoquaux, G. The NumPy Array: A Structure for Efficient Numerical Computation. Comput. Sci. Eng. 22–30 (2011).

18. Hunter, J. D. Matplotlib: A 2D graphis envinronment. Comput. Sci. Eng. 9, 90–95 (2007).

